# Canonical NLR immune receptor architecture enforces EDS1-dependency onto divergent TIR domains

**DOI:** 10.64898/2026.05.20.726525

**Authors:** Elenor Kennedy, Khong-Sam Chia, He Zhao, Jonathan DG Jones, Philip Carella

## Abstract

Toll/interleukin-1 receptor (TIR) enzymes are prominent immune components in diverse organisms across the tree of life. In flowering plants, TIRs are often integrated into nucleotide binding leucine-rich repeat (NLR) receptors whose oligomerization-dependent biochemical activities create second messengers perceived by ENHANCED DISEASE SUSCEPTIBILITY1 (EDS1)-family signaling complexes. TIR-NLRs and TIR proteins are present across the full spectrum of plant evolution, yet EDS1 signaling is a derived trait in seed plants. Here, we examined the functional dependency of diverse plant TIRs on the EDS1 pathway in the angiosperms *Nicotiana benthamiana* and *Nicotiana tabacum*. While the isolated TIR domains from non-seed plants generally required EDS1 for immune cell death activation, we also identified TIRs that functioned independent of EDS1. However, chimeric TIR-NLRs incorporating these diverse TIR domains onto the AtWRR4a receptor chassis showed a full reversion to EDS1-dependency. Extending this phenomenon further, we demonstrated that the AtWRR4a architecture enforces EDS1-dependence onto a bacterial TIR domain that is otherwise EDS1-independent. Collectively, our work demonstrates that NLR immune receptor architecture influences TIR-related immunity and provides further context to their ancient acquisition into plant immune systems.

## INTRODUCTION

Toll/interleukin-1 receptor (TIR) domains are components of immune systems across the tree of life^1,2^. To fulfill their role, they function through hydrolysis of the metabolic cofactor nicotine adenine dinucleotide (NAD^+^), small molecule production, or scaffolding^1,3,4^. In flowering plants, TIR domains are frequently integrated into nucleotide binding leucine-rich-repeat (NLR) proteins that are well-studied in crop species and are often exploited to improve disease resistance^5,6^. NLRs typically have a tripartite protein architecture, comprising a C-terminal leucine rich-repeat (LRR) domain that recognizes pathogen effectors or their perturbed host targets, a central NB-ARC domain responsible for ADP/ATP exchange during activation, and a variable N-terminal ‘signaling’ domain that biochemically initiates and/or executes downstream immune processes. NLR activation leads to robust immune responses that include mitogen-activated protein kinase (MAPK) cascades, transcriptional reprogramming, phytohormone signaling, calcium influx, and the production of reactive oxygen species^7^. Together these responses often initiate a form of programmed cell death known as the hypersensitive response (HR)^6-8^.

N-terminal domain identity is generally used to classify NLRs into functional categories. In angiosperms, NLRs contain either coiled-coil (CC) domains, RPW8-like CC domains, or TIR domains at their N-termini^8^. Once activated, many CC- and RPW8-NLRs homo-oligomerise into pentameric, hexameric^9^ or octameric^10^ structures, and their N-termini form calcium permeable toxic pores at cell membranes to initiate immune-related cell death^9^. CC-NLRs can also act as specialized ‘sensors’ that induce the oligomerization of downstream ‘helper’ NLRs that directly promote immunity^11,12^. In contrast, TIR-NLRs oligomerize to form tetrameric NADase holoenzymes that produce second messengers via the catalytic activity of a deeply conserved glutamate residue within the TIR domain^3^. To date, plant TIR domains have been shown to directly or indirectly produce a range of signaling molecules, including 2’/3’-cyclic nucleotide monophosphates (2’/3’-cNMP), 2’-cyclic ADP ribose (2’cADPR), 2ʹ-(5ʹʹ-phosphoribosyl)-5′-adenosine monophosphate/diphosphate (pRib-AMP/ADP), ADP-ribsosylated adenine triphosphate (ADPr-ATP) and ADPr-adenosine diphosphate ribose (di-ADPR), and ribofuranosyladenosine (RFA)^13-21^. The role of some of these molecules is unclear, however pRib-AMP/ADP and ADPr-ATP/di-ATP bind to and activate EDS1 (ENHANCED DISEASE SUSCEPTIBILITY1) signalling complexes that are critical for TIR-NLR-mediated immunity in angiosperms^3,19,21,22^. Both di-ADPR and ADPr-ATP bind to EDS1:SAG101 (SENESCENCE ASSOCIATED GENE 101) complexes at a C-terminal binding pocket, inducing a conformational change that promotes interaction between the EDS1 and SAG101 EP domains and the LRR domain of NRG1 (N REQUIREMENT GENE 1) family RPW8-NLRs. Binding of the EDS1:SAG101 complex is suggested to activate NRG1 by inducing conformational changes in its nucleotide binding domain, leading to activation and oligomerisation^3,17,23,24^. Similarly, pRib-AMP and pRib-ADP bind to EDS1:PAD4 (PHYTOALEXIN DEFICIENT 4) complexes at the C-terminal binding pocket. This leads to a conformational change that promotes interaction with ADR1 (ACTIVATED DISEASE RESISTANCE 1) subfamily RPW8-NLRs^3,16^. The activation of ADR1 is likely to be similar to that of NRG1, with the interfaces in PAD4 and ADR1 differing from the analogous surfaces in SAG101 and NRG1, providing specificity^24^.

Land plants evolved from freshwater charophyte algae over 500 million years ago^25^, and diverged into several lineages. These include the non-vascular seed-free bryophytes (liverworts, mosses and hornworts), vascular seed-free lineages (lycophytes and monilophytes), and seed-bearing, non-flowering gymnosperms. Angiosperms (flowering plants) diverged from non-flowering ancestors only 209 million years ago^26^. Despite being a relatively young lineage, the function and mechanisms of NLRs have primarily been studied in angiosperms, though genomic studies show that NLRs are present across land plant lineages and in some algae^27-29^. Additionally, N-terminal domains from some algal and non-flowering plant NLRs, including TIR-NLRs, have been shown to induce HR in *N. benthamiana*, suggesting a conserved immune-related function^6,30^. However, the TIR-NLR mechanism described in angiosperms is EDS1-dependent, while non-seed plants supposedly lack EDS1^6,31,32^, rendering TIR-NLR mechanism unclear in these lineages.

TIR domains are found in metazoans and bacteria, where they do not rely on EDS1 due to its absence in these lineages^32^. In bacteria, TIR enzymes are components of anti-phage immune systems like Thoeris^33,34^ and Short Prokaryotic Argonaute and associated TIR-APAZ (SPARTA)^35,36^, where they induce cell death through NAD^+^ depletion or by generating small molecule signals that activate cell death-promoting proteins^1,23,37^. Alternatively, some bacterial TIR proteins are co-opted for host-manipulation, such as the *Pseudomonas syringae* effectors HopAM1^38^ and HopBY^39^ that interfere with host immunity through NAD^+^ depletion and small molecule production^19,40^. Metazoan TIR domains often function as non-enzymatic scaffolds for protein-protein interactions, with the exception of the TIR-containing SARM1 receptor that acts as a double-stranded DNA sensor^41^ and depletes NAD^+^ to induce axonal degeneration^1,37^. The heterologous expression of metazoan and bacterial TIR enzymes in *N. benthamiana* leaves sometimes leads to EDS1-independent cell death^13,31^, indicating that TIRs can indeed function independent of the EDS1 pathway in plants.

In this study, we investigated the extent to which diverse plant TIR domains signal through the EDS1 pathway when heterologously expressed in the model flowering plants *Nicotiana benthamiana* and *Nicotiana tabacum*. We first determined that TIR domains sourced from non-flowering plants and algae show variable dependence on EDS1 through transient assays in wild-type and *eds1* mutants in *N. benthamiana*. In particular, we identified TIR domains from the moss *Physcomitrium patens* and the gymnosperm *Ginkgo biloba* that were EDS1-independent when expressed as fusions with eYFP, but become EDS1-dependent when expressed as chimeric TIR-NLRs on the AtWRR4a receptor scaffold. We further explored the influence of NLR architecture on TIR domains sourced from animals and bacteria, which revealed that the AtWRR4a NLR context enforced EDS1-dependence onto a bacterial TIR domain from the SPARTA anti-phage system. This confirmed that receptor architecture influences TIR-mediated immunity and highlights the importance of the full NLR context when interpreting TIR functional evolution.

## RESULTS

### Diverse plant TIR domains can function independently of the EDS1 pathway

TIR domains sourced from diverse plant lineages across algae-to-angiosperms (Fig 1A) activate immune-related cell death when heterologously expressed as TIR-eYFP fusions in the flowering plant *N. benthamiana*^*6*^. To determine whether these domains rely on the EDS1 signalling pathway, we expressed TIR-eYFP fusions (Fig 1B) derived from the algal species *Chlorokybus atmophyticus*, the mosses *Physcomitrium patens* and *Sphagnum fallax*, the fern *Ceratopteris richardii*, the gymnosperm *Ginkgo biloba* and the angiosperm *Arabidopsis thaliana* in wild-type (WT) and *eds1* mutant *N. benthamiana* using transient agroinfiltration assays. Alongside the angiosperm AtRPS4^TIR^ control, several TIR-eYFP fusions showed a strong dependency on EDS1 such that HR cell death was observed in WT *N. benthaminana*, but not *eds1* mutants by 7 days post-infiltration (Fig 1C; Supplementary Dataset 1). By contrast, a single TIR domain from the gymnosperm *G. biloba* (Gb.chr12.2^TIR^) and two from the moss *P. patens* (Pp3C10_21680V3^TIR^ and Pp3C22_1030V3^TIR^) were EDS1-independent, with strong cell death observed in both wild-type and *eds1* mutants. Importantly, the GUS-eYFP control failed to provoke cell death in either genotype (Fig 1C) and immunoblotting confirmed the stable expression of all tested constructs (Supplementary Dataset 2).

**Figure 1.**
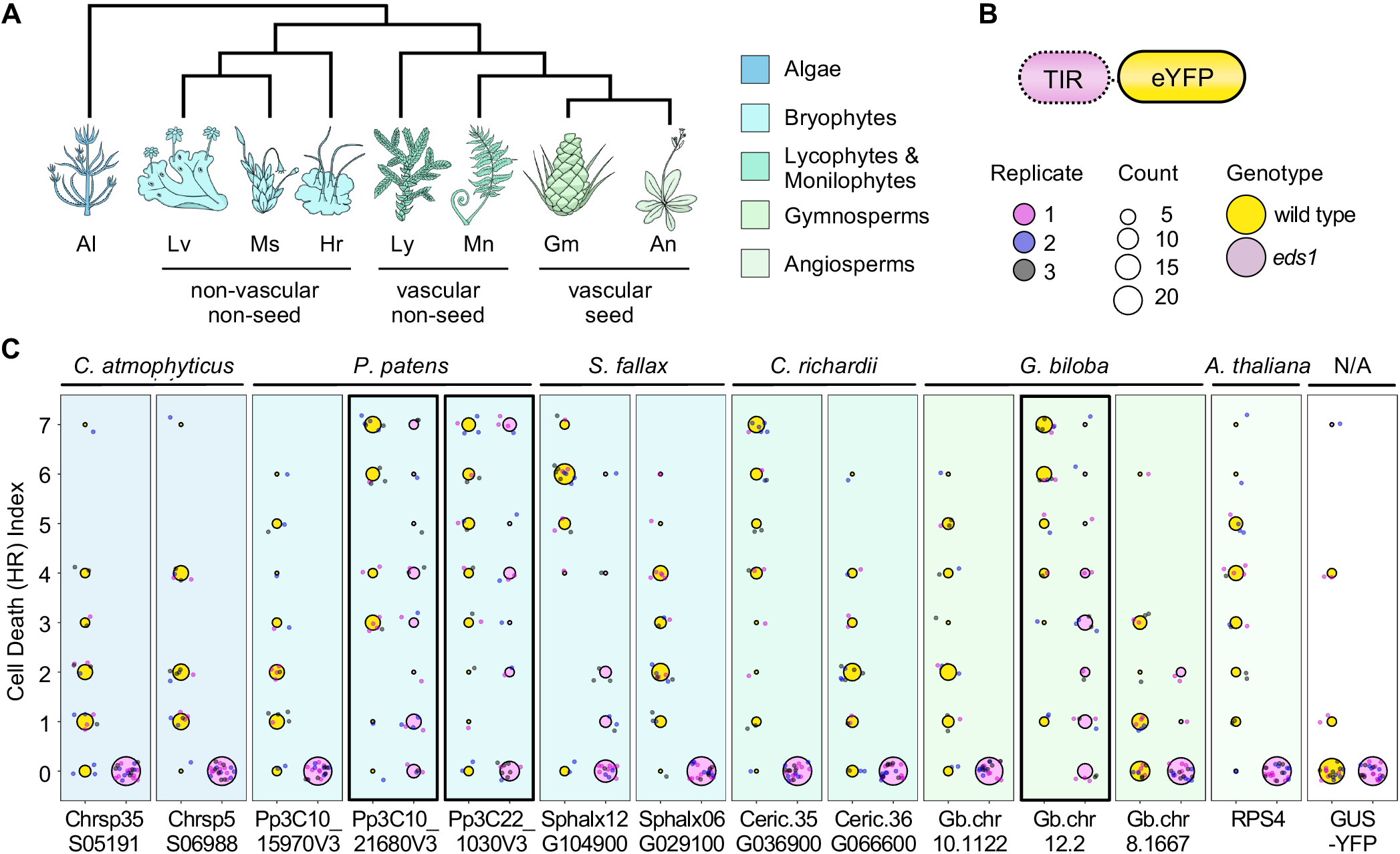
TIR-eYFP fusions from diverse plant lineages show variable EDS1-dependence. **(A)**Graphical representation of the evolutionary relationships of major land plant lineages, showing Algae (Al), Liverworts (Lv), Moss (Ms), Hornworts (Hr), Lycophytes (Ly), Monilophytes (Mn), Gymnosperms (Gy) and Angiosperms (An). Not to scale. Transitions represent a timescale of millions of years ago (mya) and are based on former estimates^25^. **(B)**Simple schematic of TIR-eYFP fusion configuration used for functional assays. **(C)** Quantification of HR cell death induced by TIR-eYFP fusions. Cell death was scored 7 days post-agroinfiltration (n ≥ 6 per replicate). Data from three independent replicates are shown in a WT and *eds1* background. Circle size indicates the number of plants with a particular score, and circle colour denotes N. benthamiana genotype. EDS1-independent domains are surrounded by a bold box.

In flowering plants, TIR-NLRs signalling through EDS1 ultimately activate the RPW8-type helpers ADR1 and NRG1 that form higher order resistosome complexes promoting immunity^9^. To assess whether non-flowering TIR domains similarly require RPW8-NLRs for their HR activity we retested all TIR-eYFP fusions in *adr1/nrg1* mutants^42^. Consistent with our previous results, we discovered that EDS1-dependent constructs elicited HR in a WT but not *adr1/nrg1* background, while the EDS1-independent constructs were able to elicit cell death in both backgrounds (Supplementary Fig. 1; Supplementary Dataset 1), further supporting their lack of dependence on the canonical EDS1-RPW8-NLR pathway. As an additional control, we tested whether diverse TIR-eYFP constructs activated HR in *nrc2/3/4* mutants lacking alternative helper CC-NLRs of the NRC network^8,43,44^. All TIR-eYFP constructs were fully functional in the *nrc2/3/4* background (Supplementary Fig. 1), ruling out the possibility that NRCs contribute to EDS1-independent cell death. Together, the data demonstrate that diverse TIRs show a general but variable reliance on the EDS1 pathway when expressed in *N. benthamiana*.

Canonical TIR-mediated responses rely on conserved catalytic glutamate residues, which are necessary for their NADase activity^3,45^ (Supplementary Figure 2). We therefore mutated each TIR domain from glutamate to alanine at this site (EA mutant) and assessed HR phenotypes in wild-type and *eds1 N. benthamiana*. As expected, all TIR-eYFP fusions with EDS1-dependent activity failed to provoke immune-related cell death when catalytic glutamates were mutated (Supplementary Fig. 3A; Supplementary Dataset 1), consistent with previous research on TIR functionality. Surprisingly, EA mutants of EDS1-independent TIRs (Gb.chr12.2, Pp3C10_21680V3, and Pp3C22_1030V3) retained HR competence in wild-type, *eds1* and *adr1/nrg1 N. benthamiana* Supplementary Fig. 3A; Supplementary Dataset 1). All constructs produced stable protein as detected by immunoblotting (Supplementary Dataset 2).

Some plant TIR domains possess 2’,3’-cAMP/cGMP synthetase activity, which is dependent upon both the conserved catalytic glutamate and a conserved cysteine residue^18^(Supplementary Figure 2). To determine whether synthetase activity contributes to the observed EDS1-independence, we created higher order cysteine to alanine mutants in the existing EA variants (CA-EA) of the EDS1-independent TIR domains from *P*.*patens* and *G*.*biloba*, and expressed them as a TIR-eYFP fusion. Similar to their WT and EA counterparts, all three CA-EA variants exhibited HR cell death in WT and *eds1* mutants (Supplementary Figure 3B; Supplementary Dataset 1), confirming that synthetase activity does not contribute to EDS1-independent responses. We confirmed expression of these TIR-eYFP fusions through immunoblotting (Supplementary Dataset 2).

### EDS1-independent TIR domains become EDS1-dependent when incorporated into chimeric TIR-NLRs

To assess the impact of the canonical TIR-NLR architecture on diverse TIR domain activity, we generated chimeric immune receptors using the well characterised *Arabidopsis* AtWRR4A receptor as a chassis^46,47^. This resulted in chimeric TIR-NLR immune receptors comprised of diverse TIR domains coupled to the NB-ARC and LRR domains of AtWRR4a (Fig. 2A). All constructs also carried a C-terminal 3x-FLAG tag to assess protein stability. Importantly, recent structural analysis demonstrates that AtWRR4a auto-inhibition is mediated only by interactions between the NB-ARC and LRR domains^46^, which we hypothesized would enable N-terminal TIR domain swaps without impacting receptor activation upon co-expression with the *Albugo candida* effector CCG28^47^ in *N. benthamiana*. Chimeric AtWRR4a receptors harboring EDS1-dependent TIR domains, including the *Arabidopsis* RPS4^TIR^-WRR4a control, all produced HR cell death in wild-type but not *eds1* mutant *N. benthamiana*, similar to the endogenous full length AtWRR4a. By contrast, AtWRR4a expressed in the absence of the CCG28 effector failed to produce an HR. Intriguingly, the EDS1-independent TIR domains (*P. patens* Pp3c10_21680V3^TIR^ and *G. biloba* Gb.chr12.2^TIR^) produced moderate HR responses in wild-type plants, but little to no response in *eds1* mutants (Figure 2B; Supplementary Dataset 1). Accumulation of TIR-WRR4a fusions was confirmed through immunoblotting (Supplementary Dataset 2), with the exception of Pp3C22_1030V3-WRR4A and Gb.chr10.1122-WRR4A constructs that induced mild EDS1-dependent cell death. This suggested that the TIR-NLR configuration limits TIR domain activity to EDS1-dependent processes. To validate this further, we generated receptor chimeras using formerly EDS1-independent TIR domains carrying catalytic EA mutations and tested for their ability to produce immune related cell death in wildtype and *eds1 N. benthamiana*. Unlike the TIR-eYFP configuration, the *P. patens* and *G. biloba* TIR-WRR4a fusions carrying EA mutations all failed to promote cell death in either background (Supplementary Fig. 3C). Collectively, the data demonstrate that the WRR4A TIR-NLR scaffold enforces EDS1-dependence and canonical catalytic activities onto diverse plant TIR domains.

**Figure 2.**
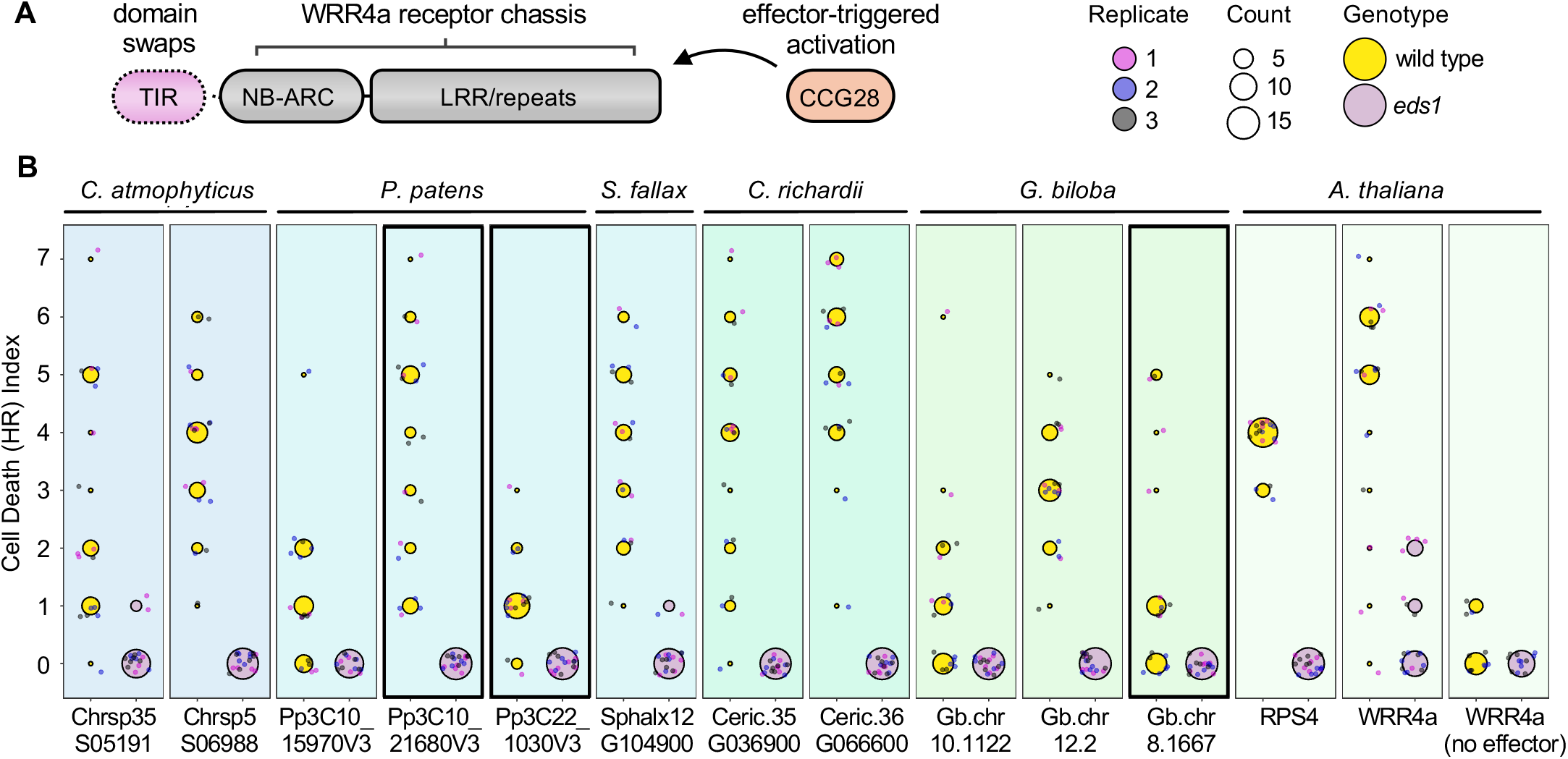
TIR-WRR4a configuration enforces EDS1-dependence and alters transcriptional response. **(A)** Schematic of TIR-WRR4a fusion. **(B)** Quantification of HR cell death induced by TIR-WRR4A fusions. Cell death was scored 7 days post-infiltration. Circle size indicates the number of plants with a particular score, and circle colour denotes *N*.*benthamiana* genotype. Data from three independent replicates are shown in a WT and *eds1* background (n ≥ 6 infiltrations per replicate). EDS1-independent domains are surrounded by a bold box. Unclustered heatmap showing differentially expressed genes (DEGs) in WT *N*.*benthamiana* when infiltrated with RPS4-YFP, RPS4-WRR4a, Pp3C10_21680V3-YFP and Pp3C10_21680V3-WRR4a, compared to a GUS-YFP control. Samples were collected 24 hours post-infiltration and variance-stabilised counts are shown.

### A canonical TIR-NLR architecture enforces EDS1-dependency onto a bacterial TIR

TIR proteins and their associated enzymatic activities are conserved across the tree of life, with similar mechanisms observed in bacteria, metazoans, and plants^1,4^. We therefore reasoned that the ability of AtWRR4a to enforce EDS1-dependency could be further extended to animal and/or bacterial TIR domains with catalytic activity. To test this, we expressed metazoan (*H. sapiens* SARM1^TIR^) and bacterial (*B. melitensis* TcpB; *M. polysiphoniae* MapTIR) TIR domains with conserved catalytic glutamate residues (Supplementary Figure 2) in both the TIR-eYFP and chimeric TIR-WRR4a configurations in wild-type and *eds1 N. benthamiana*. We observed that SARM1^TIR^-eYFP and MapTIR-eYFP induced strong EDS1-independent cell death in *N. benthamiana* (Supplementary Fig. 4A; Supplementary Dataset 1) in *N. benthamiana* leaves, whereas the RPS4^TIR^-eYFP control exhibited cell death only in wild-type plants and all other domains failed to provoke HR similar to the GUS-eYFP control. However, HR phenotypes were generally weak and variable for chimeric TIR-NLRs expressed in *N. benthamiana*, (Supplementary Fig. 4B; Supplementary Dataset 1). Protein immunoblotting confirmed accumulation of TIR-eYFP and TIR-WRR4a fusions (Supplementary Dataset 2).

We repeated this experiment in *Nicotiana tabacum*, since previous studies sometimes demonstrate stronger TIR-mediated cell death in this species^6,48,49^. Consistent with previous results, the *Physcomitrium* Pp3C10_21680V3 and *Ginkgo* Gb.chr12.2 TIR-eYFP fusions showed strong cell death in wild-type and *EDS1-*silenced *RNAi::EDS1 N. tabacum*^*50*^ (Supplementary Fig. 5A; Supplementary Dataset 1), while chimeric TIR-NLR receptors became EDS1-dependent similar to the native AtWRR4a receptor and RPS4-WRR4a (Supplementary Fig. 5B; Supplementary Dataset 1).

We then expressed SARM1^TIR^, TcpB^TIR^ and MapTIR in both TIR-eYFP and TIR-WRR4a configurations in *N. tabacum*. As anticipated, each of these domains exhibited HR phenotypes when expressed as eYFP fusions in *N. tabacum* (Fig. 3A; Supplementary Dataset 1), with SARM1^TIR^ and MapTIR exhibiting strong EDS1-independent cell death, while TcpB^TIR^ induced EDS1-dependent HR. Importantly, the RPS4^TIR^-eYFP control elicited EDS1-dependent cell death and GUS-eYFP failed to provoke an HR. All TIR-eYFP fusions were reliably detected by immunoblotting (Supplementary Dataset 2). When expressed on the AtWRR4A scaffold (Fig. 3B), SARM1^TIR^ failed to provoke an HR in both WT and *RNAi::EDS1* backgrounds and TcpB retained weak EDS1-dependent activity. By contrast, the MapTIR-WRR4a chimera provoked HR cell death in wild-type but not *RNAi::EDS1 N. tabacum* (Figure 3B; Supplementary Dataset 1). Stable expression of all receptors in *N. tabacum* was confirmed by immunoblotting (Supplementary Dataset 2). Collectively, our results demonstrate that the AtWRR4a NLR configuration can influence the activity of diverse TIR domains from plants as well as bacteria.

**Figure 3.**
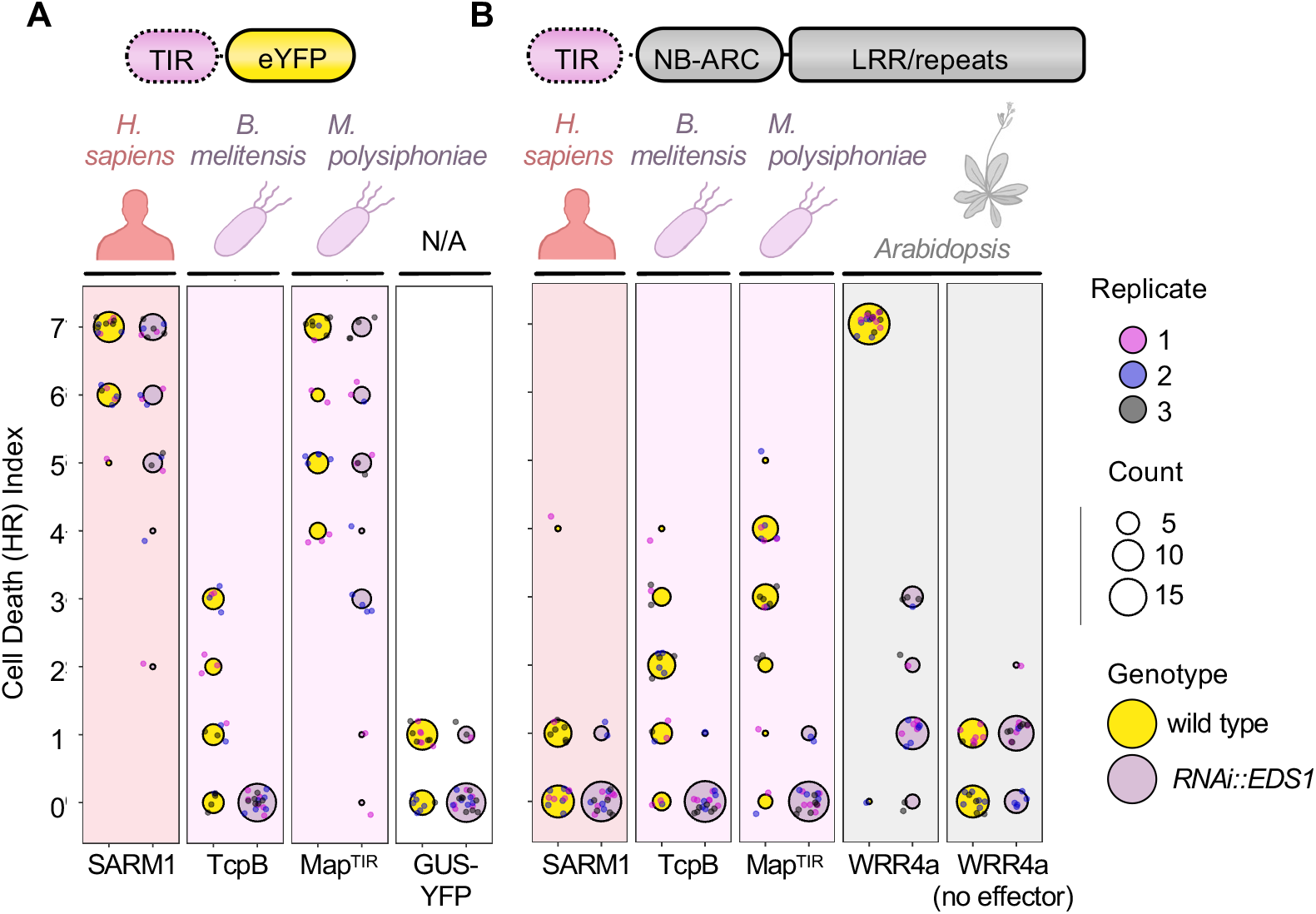
TIR-WRR4a configuration influences activity of animal and bacterial TIR domains. Quantification of cell death induced by **(A)** TIR-YFP fusions and **(B)** TIR-WRR4A fusions. Cell death was scored 7 days post-infiltration. Circle size indicates the number of plants with a particular score, and circle colour denotes *N*.*tabacum* genotype. Data from three independent replicates are shown in a WT and *RNAi::EDS1* background (n ≥ 6 infiltrations per replicate).

## DISCUSSION

In this study, we demonstrate that protein architecture impacts EDS1-dependent TIR activity. We show that the eYFP fusions routinely used to investigate TIR function^51,52^ differentially impact their reliance on EDS1 signalling relative to a canonical TIR-NLR architecture. Though many of the tested plant TIR domains remained EDS1 dependent in both the eYFP and TIR-NLR configurations, the strong alteration of EDS1 independence-to-dependence of diverse TIRs highlights the importance of protein architecture for understanding TIR functional evolution.

While we discovered a small number of EDS1-independent TIR domains, most relied on both EDS1 and the catalytic glutamate. Intriguingly, many of these TIR domains were derived from non-seed lineages that lack clear homologs of EDS1^31^. EDS1-family proteins consist of an N-terminal αβ-hydrolase (lipase-like) and a C-terminal α-helical bundle EP domain, which is thought to have arisen in the common ancestor of angiosperms and gymnosperms. While short EP-like sequences have been reported in green algae, robust EP domain homology has only been found in seed-plant lineages^32,53,54^, suggesting that this domain arose later than the TIR domain itself. Additionally, EDS1 function in TIR-NLR immunity depends upon a conserved LLIF motif in the N-terminal lipase-like domain, and a conserved FGE motif within the heterodimer cavity formed from the EP domains. These motifs are absent from non-seed plant proteins with similarity to the EDS1 N-terminal domain^32^. While non-seed lineages lack EDS1, they possess RPW8-like NLRs with shared ancestry to *Arabidopsis* ADR1-family proteins^32,55^, though it is unclear whether these are coupled to TIR domain activity as they are in Angiosperms. In any case, our data support the idea that divergent TIRs produce signals that are detected by EDS1 when expressed in *Nicotiana*. This implies that TIR activity is functionally conserved in distantly related plants, and that a functionally equivalent signaling pathway may exist in non-seed plant lineages. Future research performed directly in model non-seed plants like *P. patens* will be essential to understand both the chemical messengers produced by diverse TIR-NLRs and the unique signalling pathways used to transduce them.

TIR domains from non-flowering *P. patens* and *G. biloba* promoted HR cell death independent of both EDS1 and the catalytic glutamate, a conserved residue that is essential for both NADase and nucleic acid metabolism in other plant TIRs^3,18,45^. Prior knowledge of EDS1-independent TIR domains is largely limited to non-plant enzymes thought to function primarily through NAD^+^-depletion and potentially via NAD-derived compounds^19,40^, a phenomenon that relies on the conserved glutamate for activity. In plants, a TIR domain from the *Zea mays* TNP receptor (TIR-NBARC-TPR configuration) was found to be similarly EDS1-independent when expressed in *N*.*tabacum*, however, its activity was dependent on the conserved catalytic glutamate residue^31^. Given that the *P. patens* and *G. biloba* EDS1-independent TIR domains did not depend on this residue, they likely function via an alternative mechanism. The *Arabidopsis* TIR-NLR RBA1 and the flax (*Linum usitatissimum*) TIR-NLR L7 have been proposed to have 2’,3’-cAMP/cGMP synthetase activity that relies on the conserved catalytic glutamate and a conserved cysteine residue^3,18^. This too is unlikely to account for the EDS1-independent activity of *P. patens* and *G. biloba* TIRs, as mutating both residues had no impact on their ability to provoke cell death. Collectively, these data indicate that the *P. patens* and *G. biloba* TIR-eYFP fusions function through an unknown, non-catalytic mechanism to provoke cell death. Importantly, these TIR domains retain EDS1-dependent activity when expressed in a TIR-NLR configuration, indicating functional conservation that was otherwise masked when overexpressed on the YFP scaffold.

Prokaryotic TIR domains like HopAM1 and TcpO induce EDS1-independent cell death through NAD^+^ depletion and/or by producing small molecules that do not utilize the EDS1 pathway^19,40^. Intriguingly, some prokaryotic TIR domains do produce NAD^+^-derived molecules similar to those of plant TIRs that signal via EDS1. For example, TcpB/BtpA from *Brucella melitensis* produces 2’cADPR^2^, a compound produced by plant TIR domains^2,39,56-58^. The *Acinetobacter baumanii* TIR (AbTIR) and *Pseudomonas* HopBY TIR proteins also produce 2’cADPR that can induce EDS1-PAD4-ADR1 complex formation when expressed in insect cells, and induce cell death via this complex in *N. benthamiana*^*19*^. Our work identified EDS1-dependent activies for TcpB (in any configuration) and MapTIR (in an NLR configuration) that further support the idea that bacterial TIRs can function through EDS1. Exactly how MapTIR activity is influenced by its overall protein architecture remains to be determined. We hypothesize that the TIR-eYFP context is permissive to alternative mechanisms, whereas the TIR-NLR configuration likely constrains TIR activities to produce canonical signaling molecules detected by EDS1. Whether 2’cADPR is essential for canonical EDS1-signalling also remains to be determined, as MapTIR lacks a conserved tryptophan residue required for ADPR cyclisation^58^. Further structural and biophysical data are required to understand the impact of NLR architecture on TIR enzymatic activity.

Metazoan TIR domains are thought to function as scaffolds rather than NADase enzymes. Toll-like receptors (TLRs) and interleukin-1 receptors (IL-1Rs) activated by pathogen- or -damage associated molecular patterns (PAMPs/DAMPs) dimerize and recruit cytoplasmic adaptor proteins like MyD88, which also contain TIR domains. These adaptors activate downstream signaling proteins, inducing inflammatory responses and cell death^1^. By contrast, the metazoan SARM1 receptor functions as an octameric NADase that promotes axonal degeneration through NAD^+^ depletion^1,37^. SARM1^TIR^ has previously been demonstrated to induce EDS1-indendent cell death in *N. benthamiana*, likely due to NAD^+^ depletion, which is not observed upon expression of plant TIRs^13,21^. Intriguingly, a recent study shows that metazoan ‘scaffolding’ TIRs like the TLR4^TIR^ produce NAD^+^-derived cADPR when expressed *in vitro*, hinting towards greater functional diversity across animals^59^. Future studies addressing the biochemical diversity of TIR-derived products and their activities across the tree of life will advance our understanding of these key immune proteins.

TIR domain-containing proteins typically require a stimulus to promote oligomerization that enables their enzymatic activity. In the absence of known stimuli, TIR domain functionality has been tested by generating translational fusions to different scaffolds that permit TIR-association and enzymatic activity. Overexpression of TIR domains fused to YFP or other small epitope tags, as performed in our study, has been shown to permit the ‘autoactivation’ of many plant TIR domains, including flax (*Linum usitatissimum)* L6^52,60,61^, *Arabidopsis thaliana* RPS4^6,61^, and diverse TIRs from algae and non-flowering plants^6,30^. Induced proximity using alternative fusions has also been used to examine TIR function. For example, fusions incorporating plant TIR domains onto the mammalian NLRC4 receptor enabled TIR-mediated HR through induced oligomerisation induced by mammalian NAIP proteins and pathogen-associated molecular patterns^50^. Notably, both SARM1^TIR^ and AbTIR were incapable of inducing HR in *N*.*benthamiana* on the NLRC4 scaffold^50^, despite inducing EDS1-independent and glutamate-dependent cell death when expressed alone^40^, supporting the idea that different scaffolds can alter TIR domain activity. SARM1^TIR^ was similarly inactive in the chimeric AtWRR4a context, which is likely due to improper assembly of TIR oligomers since SARM1 is natively octameric^62^ rather than tetrameric.

Our work demonstrates that NLR architecture influences the EDS1-dependence of diverse TIR domains. This highlights the importance of considering native protein architectures when examining the activity of TIR enzymes, which likely applies to the full diversity NLR N-terminal ‘signaling’ domains. For TIR-NLRs, future research should consider the full receptor configuration, using known effectors and/or ‘autoactive’ MHD^52^ mutants where possible, or at least by adopting a chimeric activation system like the AtWRR4a receptor and CCG28 effector system presented here. Indeed, a recent examination of Arabidopsis TIR activity leveraged a chimeric TIR-NLR chassis utilizing the Arabidopsis RPP1 receptor and ATR1 effector to reveal consensus cell-death activation motifs^21^. Our results on divergent TIRs and contrasting protein architectures supports the idea that TIR-NLR activity is conserved in land plants and hint towards commonalities in TIR-mediated immunity in diverse plant lineages that lack EDS1. Future research aimed at resolving TIR-NLR evolution in streptophyte algae and bryophytes is needed to pinpoint the precise acquisition and co-option of this receptor strategy, which may have evolved early to further constrain TIR domain activity for immune-related purposes.

## ACKNOWLEDGEMENTS

We thank the TSL SynBio platform for access to the cloning vectors used in this study. We also thank all current and past members of the Carella group at the John Innes Centre for technical assistance and insightful discussions on TIR functional evolution. We thank Henriette Laessle (JIC) for helpful discussions on TIR enzymatic activities.

## AUTHOR CONTRIBUTIONS

E.B.K., K-S.C., H.Z, J.D.G.J, and P.C. designed the research; E.B.K., and K-S.C. performed the research; E.B.K. and K-S.C. analyzed the data; E.B.K. and P.C. prepared figures and final datasets; E.B.K and P.C. wrote the manuscript with contributions from all authors.

## DATA AVAILABILITY

All relevant gene identifiers are presented in the manuscript and in the supporting information.

## FIGURE LEGENDS

**Supplementary Figure 1.**
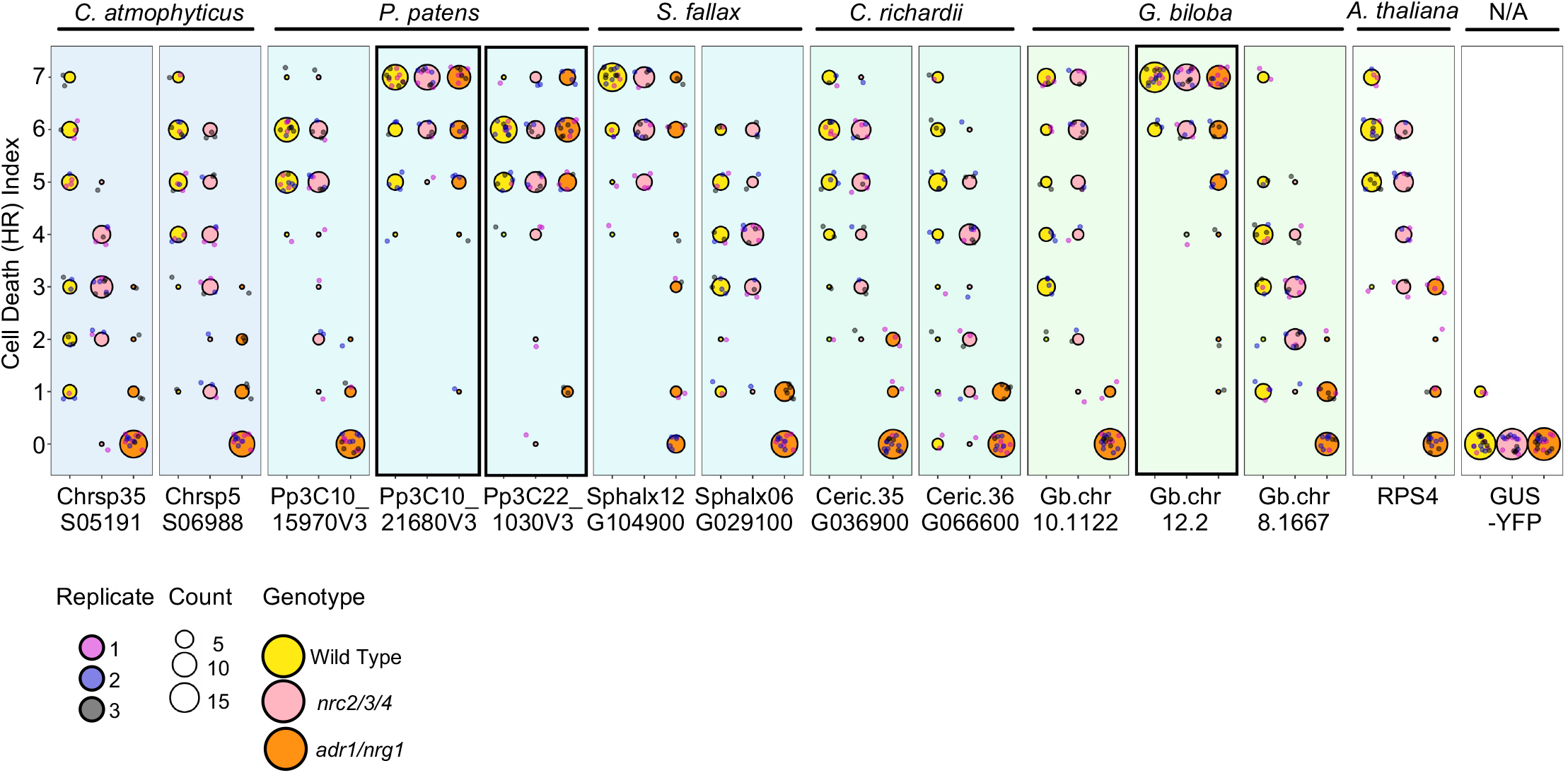
Quantification of HR cell death induced by TIR-eYFP fusions. Cell death was scored 7 days post-infiltration (n ≥ 6 infiltrations per replicate). Circle size indicates the number of plants with a particular score, and circle colour denotes *N*.*benthamiana* genotype. Data from three independent replicates are shown in a WT, *nrc/2/3/4* and *adr1/nrg1* background. EDS1-independent domains are surrounded by a bold box.

**Supplementary Figure 2.**
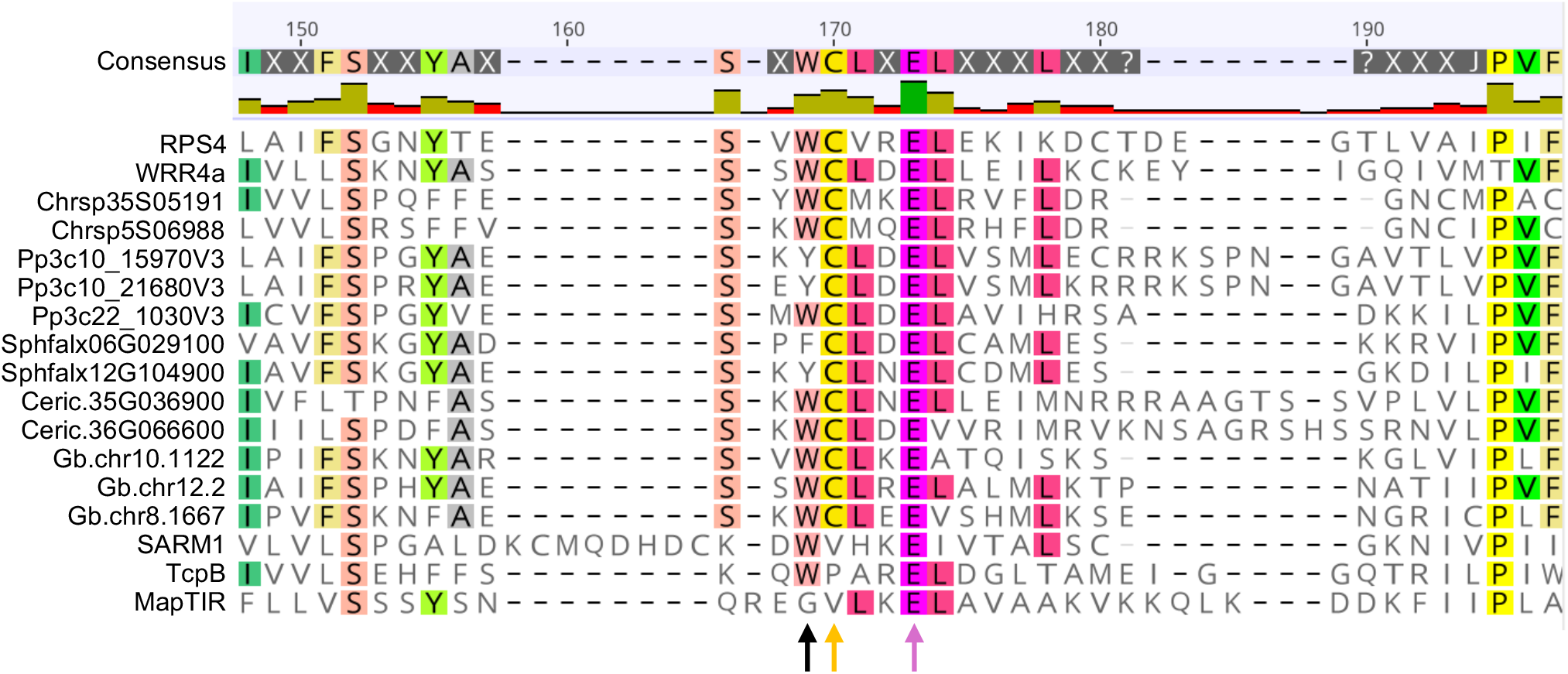
Alignment of conserved catalytic glutamate (indicated by black arrow) required for NADase activity. The position of the conserved cysteine required for 2’,3’-cAMP/cGMP synthetase activity is indicated by a green arrow. TIR domains were aligned using MUSCLE alignment in Geneious Prime (2026.0.2). The consensus sequence represents residues present in over 50% of the sequences. Default colours were used, and are indicative of the residues’ physicochemical properties. Only residues matching the consensus sequence are coloured.

**Supplementary Figure 3.**
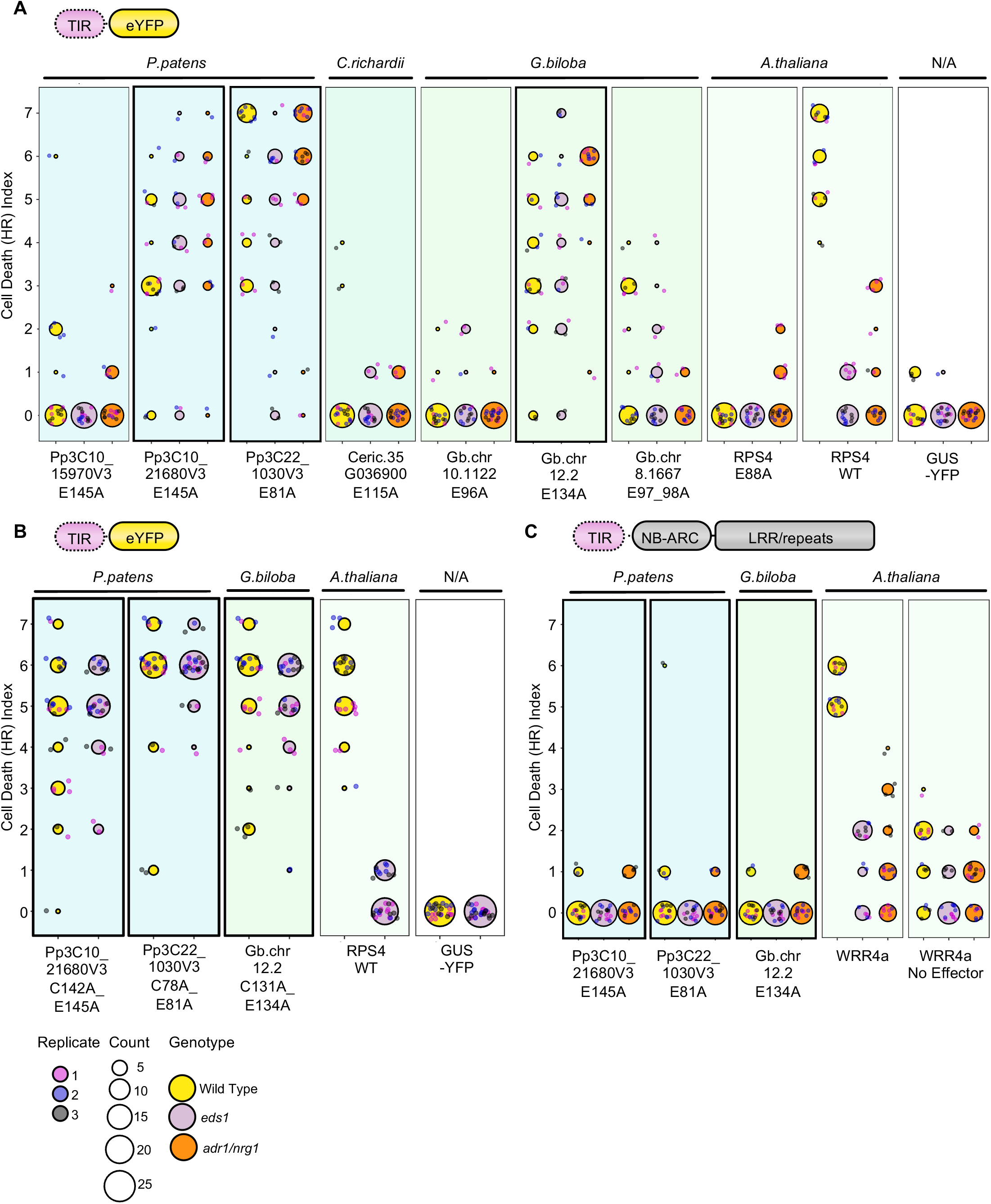
Quantification of HR cell death induced by **(A)** TIR-eYFP fusions in which the catalytic glutamate required for NADase activity is mutated to alanine in WT, *eds1* and adr1/nrg1 *N*.*benthamiana*, **(B)** TIR-eYFP fusions in which a conserved cysteine residue required for 2’,3’-cAMP/cGMP synthetase activity is mutated to alanine in WT and *eds1 N*.*benthamiana* and **(C)** TIR-WRR4a fusions in which the catalytic glutamate is mutated to alanine in WT, *eds1* and *adr1/nrg1 N*.*benthamiana*. Cell death was scored 7 days post-infiltration. Circle size indicates the number of plants with a particular score, and circle colour denotes *N*.*benthamiana* genotype. Data from three independent replicates are shown in a WT and *eds1* background.

**Supplementary Figure 4.**
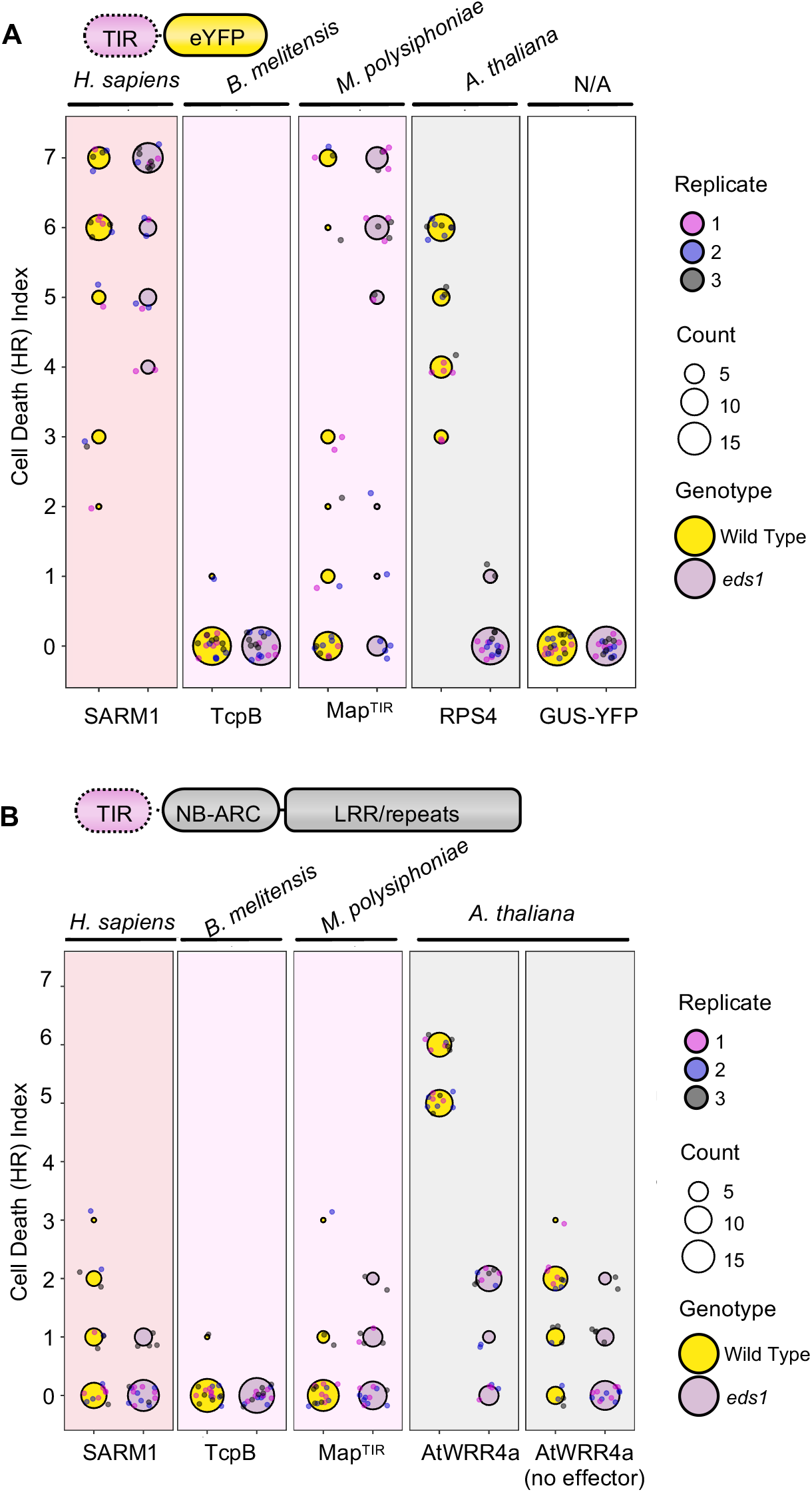
Quantification of cell death induced by **(A)** TIR-YFP fusions and **(B)** TIR-WRR4A fusions. Cell death was scored 7 days post-infiltration. Circle size indicates the number of plants with a particular score, and circle colour denotes *N. benthamiana* genotype. Data from three independent replicates are shown in a WT and *eds1* background.

**Supplementary Figure 5.**
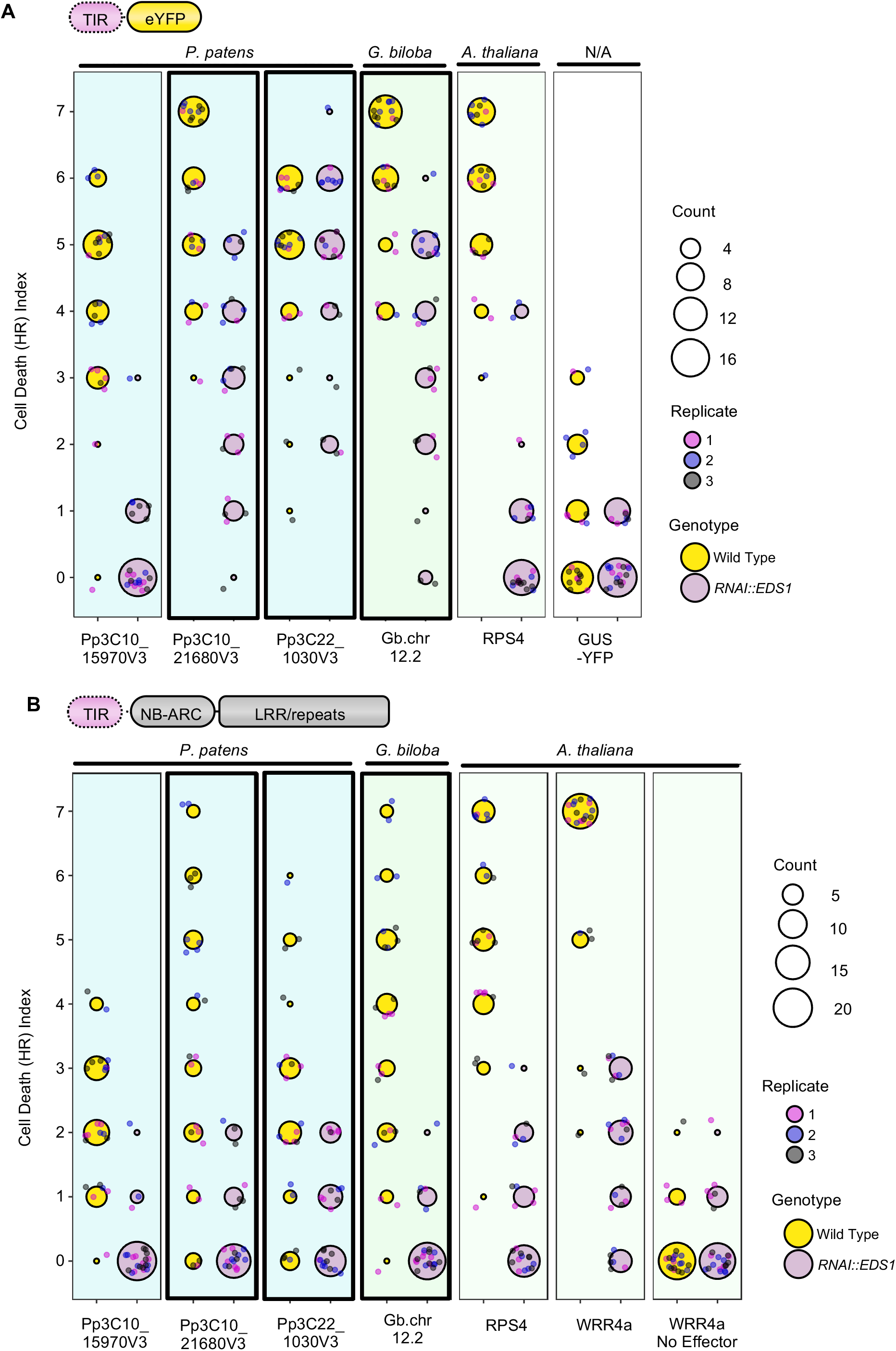
Quantification of cell death induced by **(A)** TIR-YFP fusions and **(B)** TIR-WRR4A fusions. Cell death was scored 7 days post-infiltration. Circle size indicates the number of plants with a particular score, and circle colour denotes N.tabacum genotype. Data from three independent replicates are shown in a WT and *RNAi::EDS1* background. EDS1-independent domains are surrounded by a bold box.

## SUPPLEMENTARY INFORMATION

### Supplementary Figures

**Supplementary Dataset 1. Representative images of HR cell death**.

**Supplementary Dataset 2. Protein immunoblots**

**Supplementary Dataset 3. Genomic resources and construct sequecences**.

## MATERIALS AND METHODS

### Plant and Bacterial Growth Details

*Nicotiana benthamiana* and *Nicotiana tabacum* ‘Samsun’ plants were grown in a controlled environment, in soil, at 22 degrees Celsius, with a 16-hour photoperiod (160-200uE, fluorescent, Sylvania F58W/GRO). *Escherichia coli* NEB 5-α cells were cultured overnight in LB medium supplemented with appropriate antibiotics at 37 °C with shaking. *Agrobacterium tumefaciens* GV3103 carrying pMP90 cells were cultured in LB medium supplemented with appropriate antibiotics at 28 °C overnight, with shaking.

### Cloning

TIR domains, flanked by BsaI restriction sites, were synthesised as described by Chia *et al*.,2024^6^. All TIR domains and constructs used can be found in Supplementary Dataset 3. TIR-WRR4A fusions consisted of a TIR domain of interest and the C-terminus of AtWRR4A-3xFLAG^47^ from the start of the NB-ARC domain onwards. Primers used in this study are available in Supplementary Dataset 3. These were assembled in a Golden Gate compatible system with pICH85281 (mannopine synthase promoter + W, Addgene #50332) and pICH41276 (*Arabidopsis* heat shock protein 18 terminator, TSL, SynBio) in the binary vector pICH47742 (Addgene #48001). Golden Gate products were transformed into NEB 5-α *Escherichia coli* cells (New England Biolabs) using a heat shock method. Bacteria were plated on LB media containing carbenicillin, and the presence of the gene of interest in surviving white colonies was confirmed through colony PCR (primers used are in Supplementary Dataset 3). The resultant plasmid was purified using the QIA Spin Miniprep Kit (Qiagen, cat no. 27104) according to manufacturer guidelines. Plasmids were then sequenced using Sanger sequencing (Genewiz, Azenta Life Sciences).

Sequenced plasmids were transformed into *Agrobacterium tumefaciens* GV3103 cells by electroporation. Resultant transformed bacteria were grown on LB media containing Carbenicillin, Rifampicin and Gentamycin for selection. Transformation success was checked using colony PCR. PCR primers used are in Supplementary Dataset 3.

### Agroinfiltration Cell Death Assays

Cell death assays were performed as described in Chia *et al*., 2024^6^. In brief, the expanded leaves of 4-week-old *N. benthamiana* or *N. tabacum* plants were infiltrated with *Agrobacterium tumefaciens* carrying a binary of expression vector for the TIR domain of interest. Bacterial suspensions were prepared in fresh inoculation buffer (10mM MES-KOH, 10mM MgCl_2_, 150mM acetosyringone, pH5.6). Suspensions were adjusted to OD_600_ = 0.3. In *N. benthamiana*, bacteria expressing the TIR domain of interest were mixed in a 1:1 ratio with *Agrobacterium* expressing the silencing suppressor P19. P19 was not included for *N*.*tabacum* where it triggers cell death. For WRR4A fusions, *Agrobacterium* were mixed in a 1:1 ratio with bacteria expressing the effector CCG28^47^ for activation. Leaves were harvested 7 days post-inoculation and the resultant cell death phenotype was scored on a scale of 0-7, between no visible cell death and fully confluent cell death. Representative cell death pictures can be found in Supplementary Dataset 1. For YFP fusions, GUS-YFP was used as a negative control. For WRR4A fusions, AtWRR4A with CCG28 was used as a positive control, and AtWRR4A with no effector was used as a negative control. Cell death graphs were plotted using ggplot2 (https://cran.r-project.org/web/packages/ggplot2/citation.html) in R^63^.

### Immunoblotting

Protein samples were prepared using 6 x 4 mm tissue discs, sampled from *N. benthamiana* or *N. tabacum* leaves. Samples were taken 3 days post-inoculation, unless they caused strong cell death, in which case they were sampled at 3 days post-inoculation. Samples were frozen in liquid nitrogen and ground, then stored at -80 °C before use. Samples were homogenised in 2X LDS solution, prepared using mPAGE™ 4X Sample Buffer (Merck Millipore, cat no. MPSB-10ML), according to manufacturer instructions.

Plant-derived TIR-eYFP fusions in *N. benthamiana* were run on an 4-12% NuPAGE− Bis-Tris Mini Protein Gel (Invitrogen, NP0321) with MOPS buffer. Animal and bacterial TIR-eYFP and TIR-WRR4a constructs in *N*.*benthamiana* were run on 12% NuPAGE− Bis-Tris Mini Protein Gel (Invitrogen, NP0341). Remaining constructs were run in an 8% Bis-Tris gel with MOPS buffer. For YFP fusions, immunoblotting was performed with a primary anti-GFP antibody (11814460001, Roche, 1:2500 dilution) and secondary HRP-linked Anti-Mouse IgG (NXA931-1ML, Amersham, 1:5000 dilution), or using using HRP-conjugated anti-GFP antibody (Miltenyi Biotech, 130-091-833, 1:5000 dilution). For WRR4A fusions, immunoblotting was performed either with mouse-derived monoclonal ANTI-FLAG® M2-peroxidase (HRP) antibody (Sigma-Aldrich, A8592), or using primary ANTI-FLAG® M2 antibody, Mouse monoclonal (Sigma-Aldrich, F1804), followed by secondary HRP-linked Anti-Mouse IgG (NXA931-1ML, Amersham, 1:5000 dilution). Total protein loading was visualised on nitrocellulose membranes through staining with Ponceau S solution (Sigma-Aldrich, P7170).

### Bioinformatics

Plant TIR proteins were initially identified using NLRTracker^64^, as described in Chia *et al*., 2024^6^. This was applied to the relevant genome annotation for each species (*Chlorokybus atmophyticus* – CCAC 022011, *Spaghnum fallax* – v1.1 (http://phytozome.jgi.doe.gov/), *Physcomitrium patens* – v3.312, *Ceratopteris richardii* – v2.113 (Marchant *et al*., 2024), *Gingko biloba*^*65*^, *Arabidopsis thaliana* – Araport11 (https://phytozome-next.jgi.doe.gov/)). The sequence for MapTIR was obtained from Koopal *et al*., 2022^35^. The sequence of all other bacterial and animal TIR domains were obtained from the PDB via Nimma *et al*., 2021^1^. All genomic resources and TIR construct sequences are available in Supplementary Dataset 3.

