## Supplementary Data 1 for "Canonical NLR immune receptor architecture enforces EDS1-dependency onto divergent TIR domains"

### A. TIR-eYFP Fusions in Wild Type *N. benthamiana*

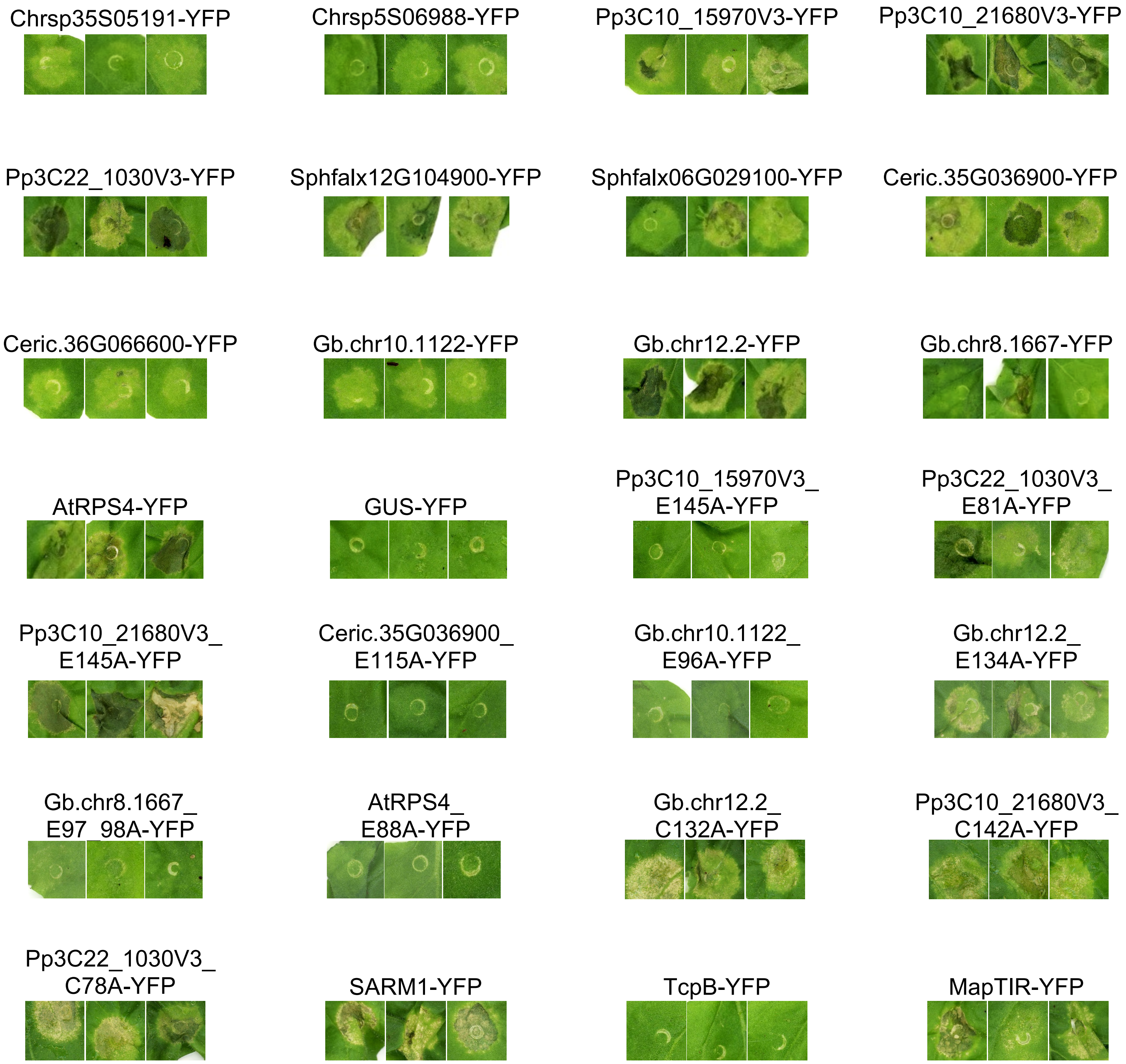


**Supplementary Dataset 2. Macroscopic Cell Death Phenotypes.**
**(A)** Representative cell death phenotypes for TIR-eYFP fusions in wild-type *N. benthamiana*, photographed 7 days post-infiltration and cropped from independent leaves.

#### B. TIR-eYFP Fusions in *eds1 N. benthamiana*

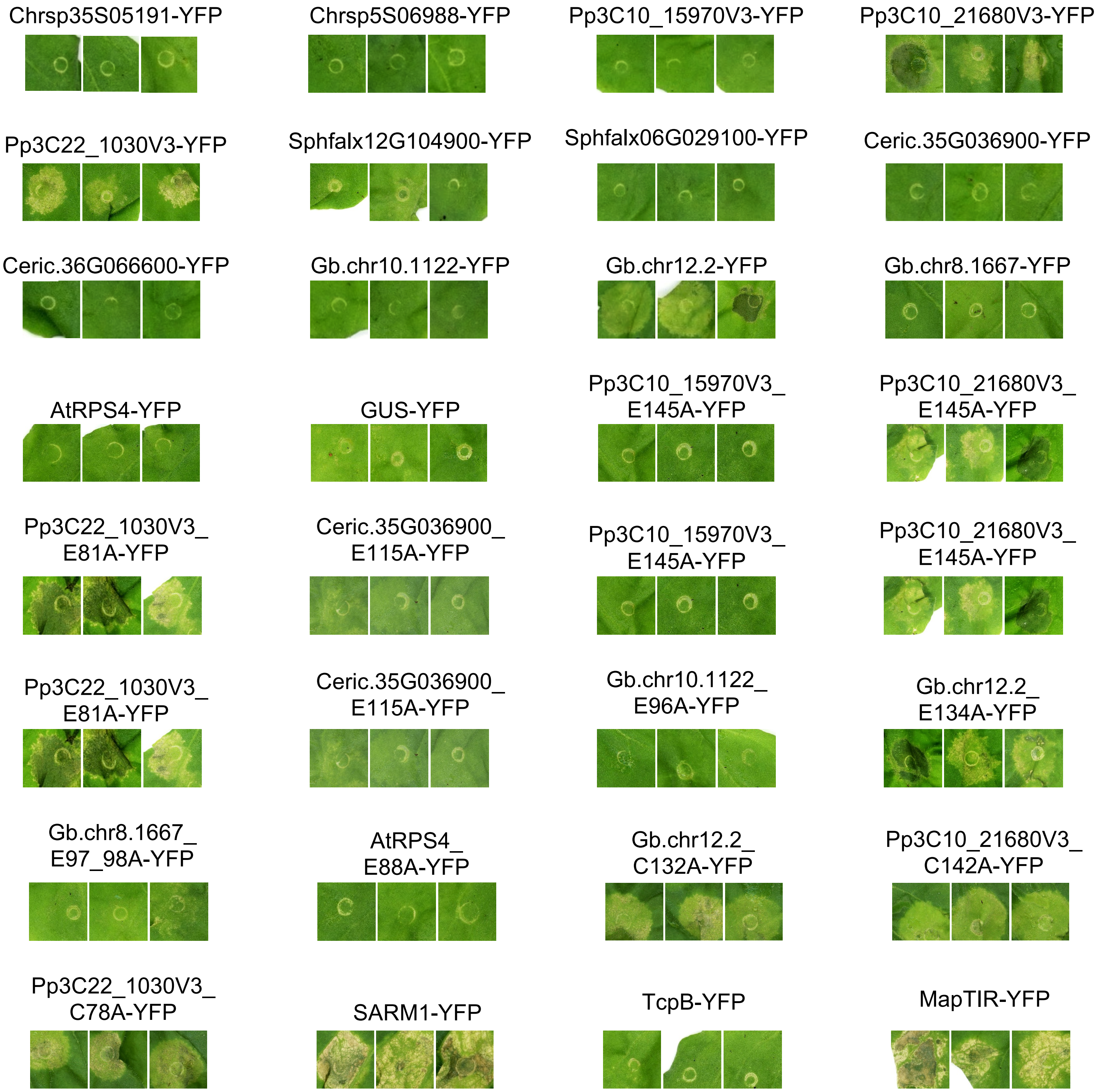


**Supplementary Dataset 2. Macroscopic Cell Death Phenotypes.**
**(B)** Representative cell death phenotypes for TIR-eYFP fusions in *eds1 N. benthamiana*, photographed 7 days post-infiltration, and cropped from independent leaves.

#### C. TIR-eYFP Fusions in *nrc2/3/4 N.benthamiana*

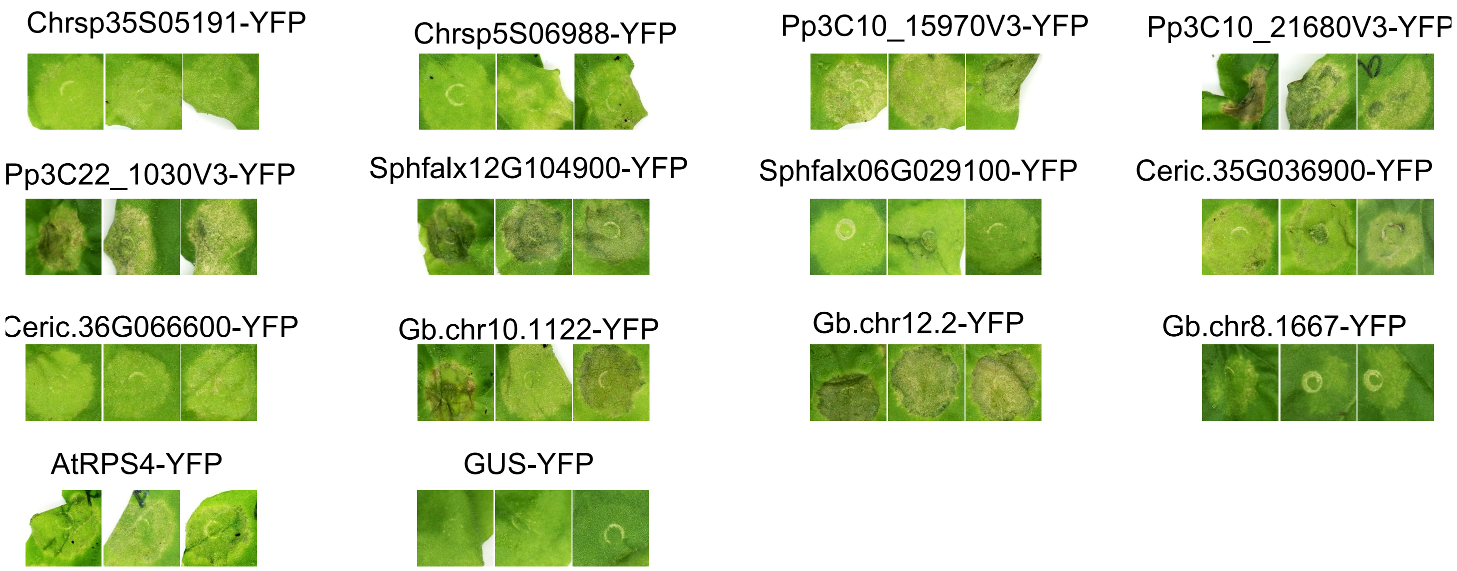


#### D. TIR-eYFP Fusions in *adr1/nrg1 N.benthamiana*

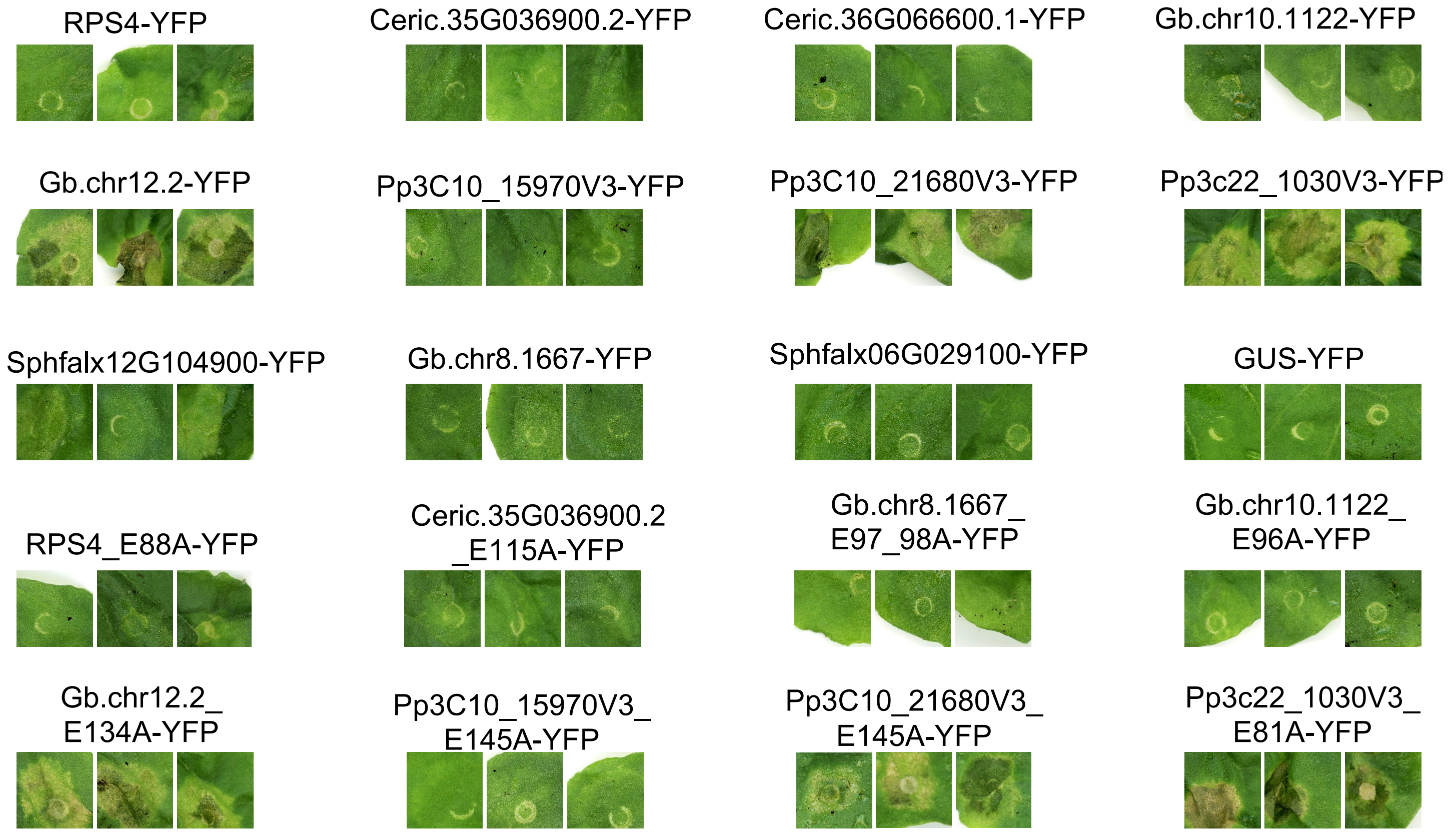


**Supplementary Dataset 2. Macroscopic Cell Death Phenotypes.**
**(C)** Representative cell death phenotypes for TIR-eYFP fusions in *nrc2/3/4 N. benthamiana*, photographed 7 days post-infiltration, and cropped from independent leaves.
**(D)** Representative cell death phenotypes for TIR-eYFP fusions in *adr1/nrg1 N. benthamiana*, photographed 7 days post-infiltration, and cropped from independent leaves.

#### E. TIR-WRR4A Fusions in Wild Type *N.benthamiana*

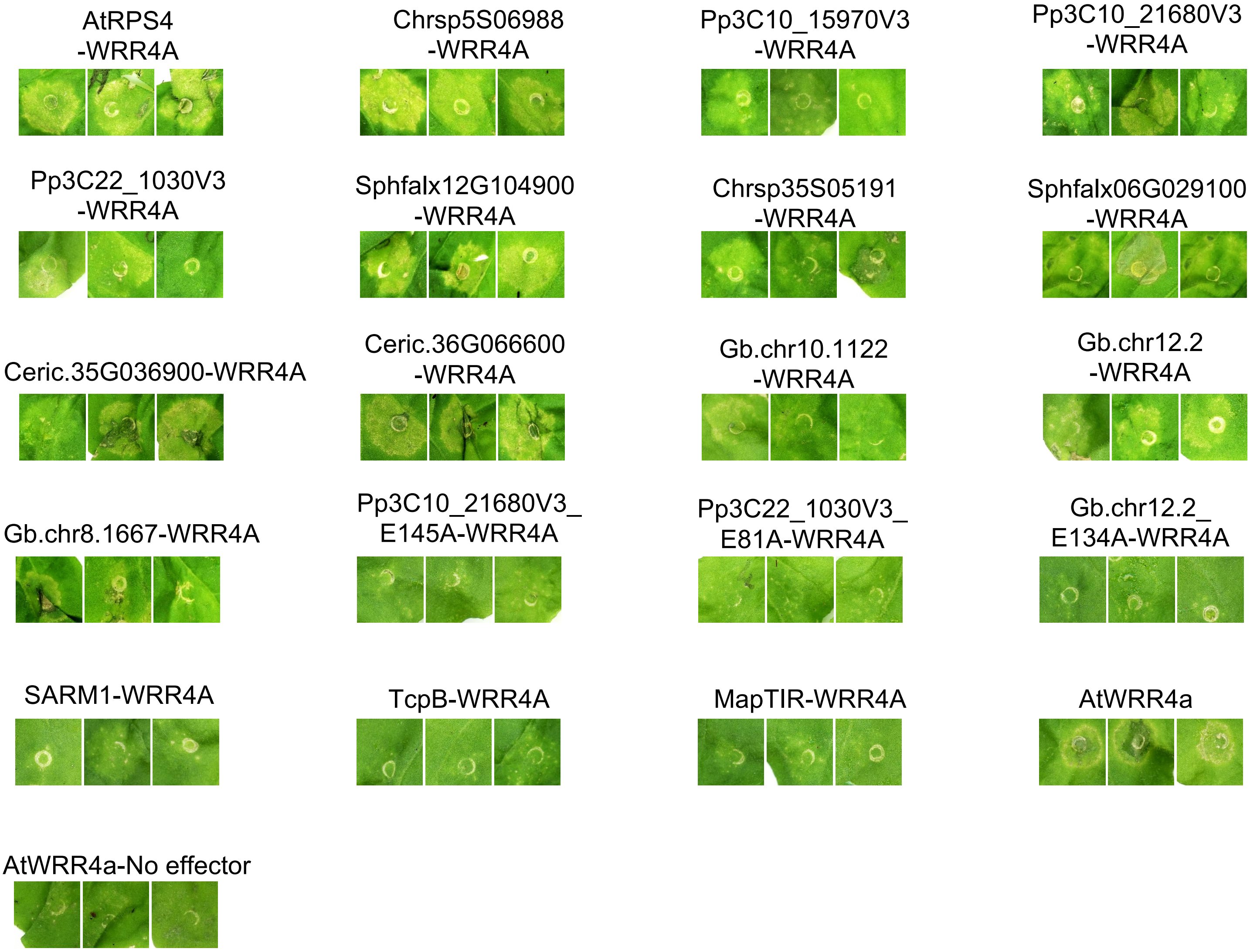


**Supplementary Dataset 2. Macroscopic Cell Death Phenotypes.**
**(E)** Representative cell death phenotypes for TIR-WRR4a fusions in wild-type *N. benthamiana*, photographed 7 days post-infiltration, and cropped from independent leaves.

#### F. TIR-WRR4A Fusions in *eds1 N.benthamiana*

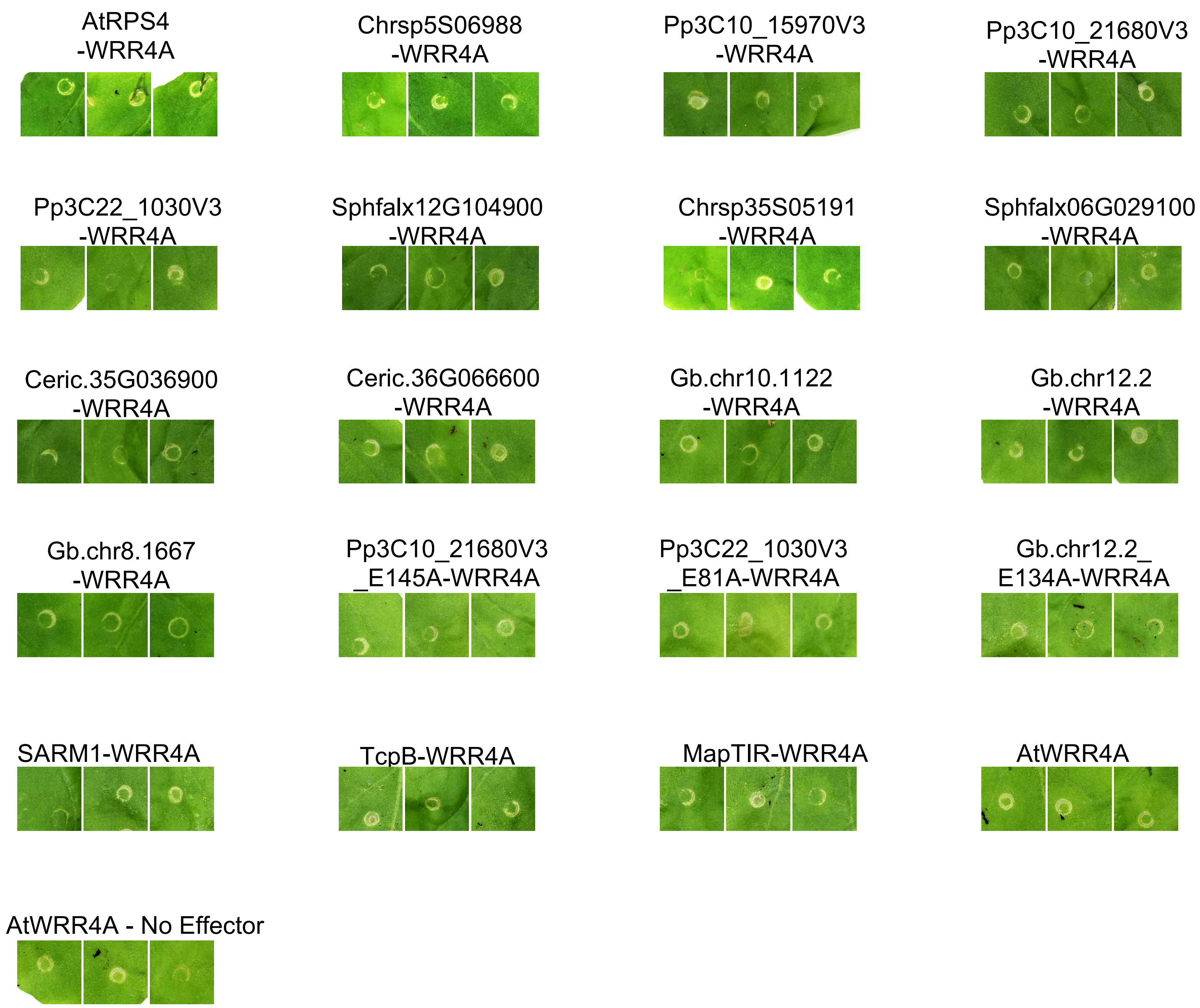


#### G. TIR-WRR4a Fusions in *adr1/nrg1 N.benthamiana*

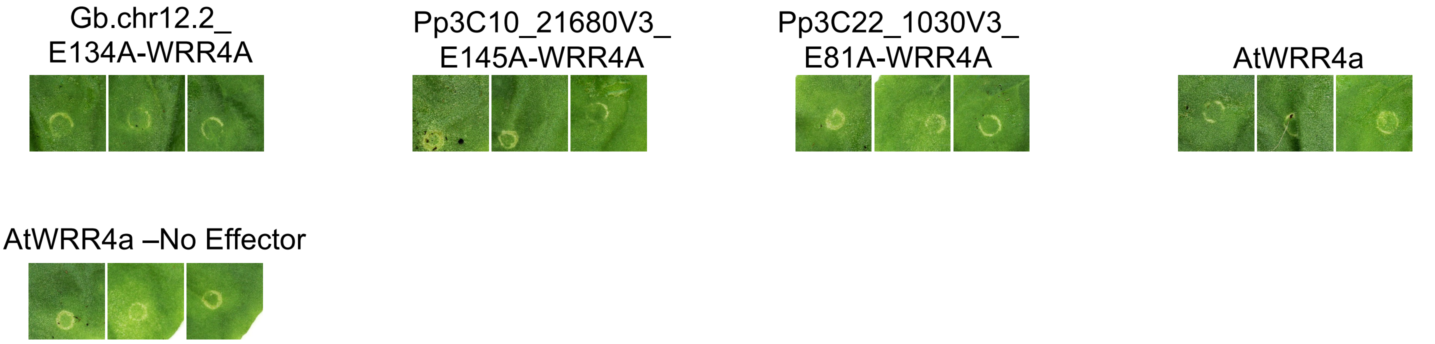


**Supplementary Dataset 2. Macroscopic Cell Death Phenotypes.**
**(F)** Representative cell death phenotypes for TIR-WRR4a fusions in *eds1 N. benthamiana*, photographed 7 days post-infiltration, and cropped from independent leaves.
**(G)** Representative cell death phenotypes for TIR-WRR4a fusions in *adr1/nrg1 N. benthamiana*, photographed 7 days post-infiltration, and cropped from independent leaves.

#### H. TIR-eYFP Fusions in Wild Type *N.tabacum*

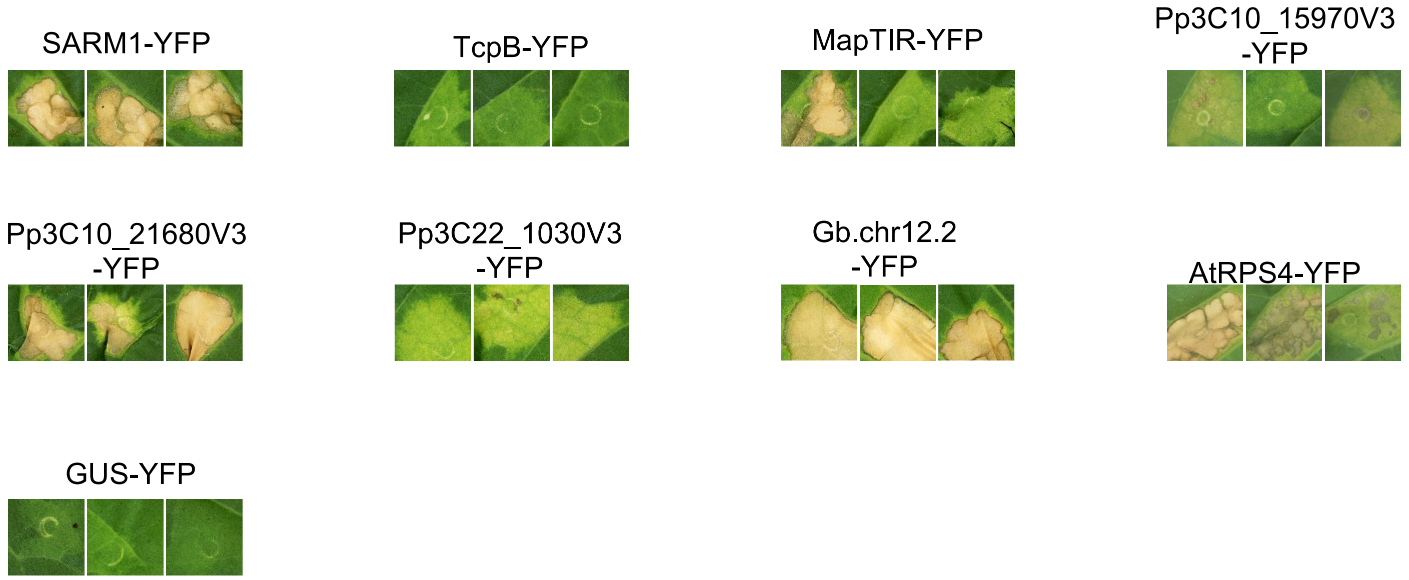


#### I. TIR-eYFP Fusions in *RNAi:: EDS1 N.tabacum*

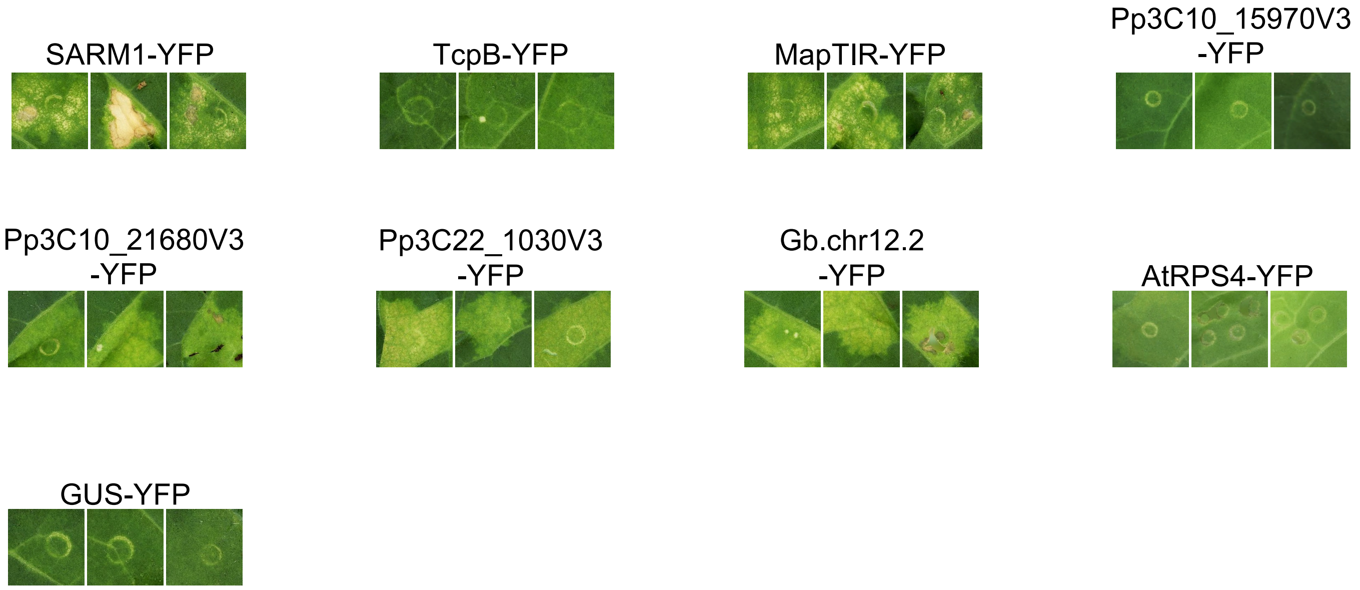


**Supplementary Dataset 2. Macroscopic Cell Death Phenotypes.**
**(H)** Representative cell death phenotypes for TIR-eYFP fusions in *eds1 N. tabacum*, photographed 7 days post-infiltration, and cropped from independent leaves.
**(I)** Representative cell death phenotypes for TIR-eYFP fusions in *RNAi::EDS1 N. tabacum*, photographed 7 days post-infiltration, and cropped from independent leaves.

#### J. TIR-WRR4A Fusions in Wild Type *N.tabacum*

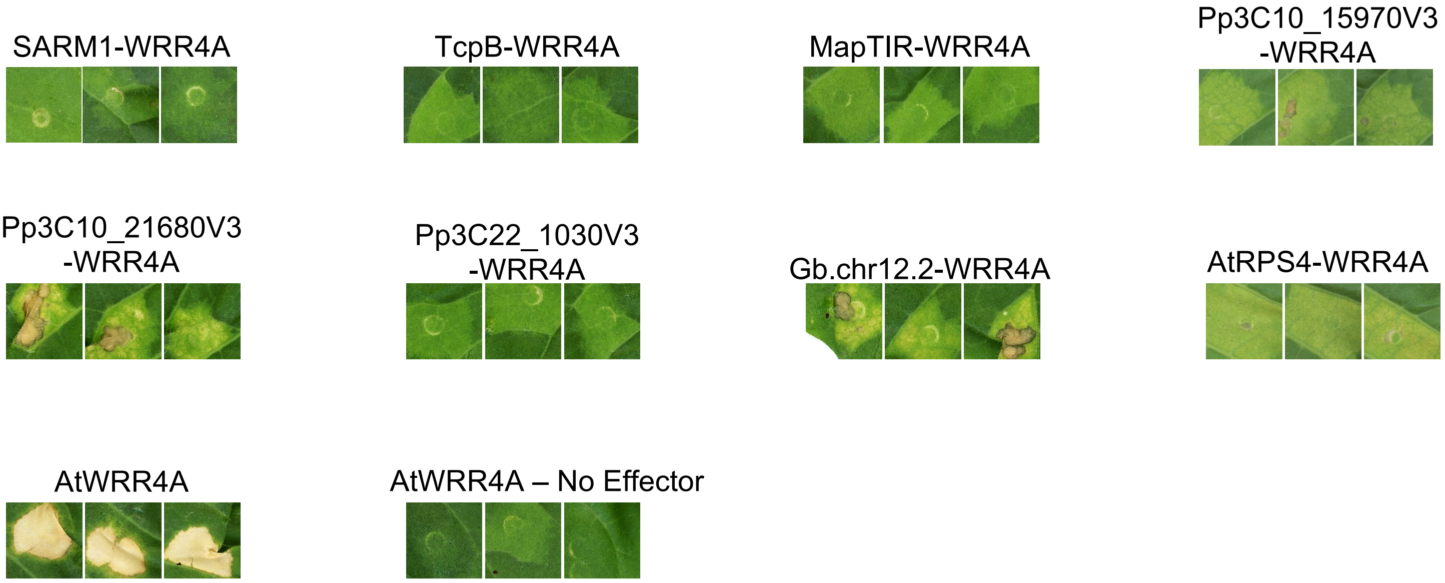


#### K. TIR-WRR4A Fusions in RNAi::EDS1 *N.tabacum*

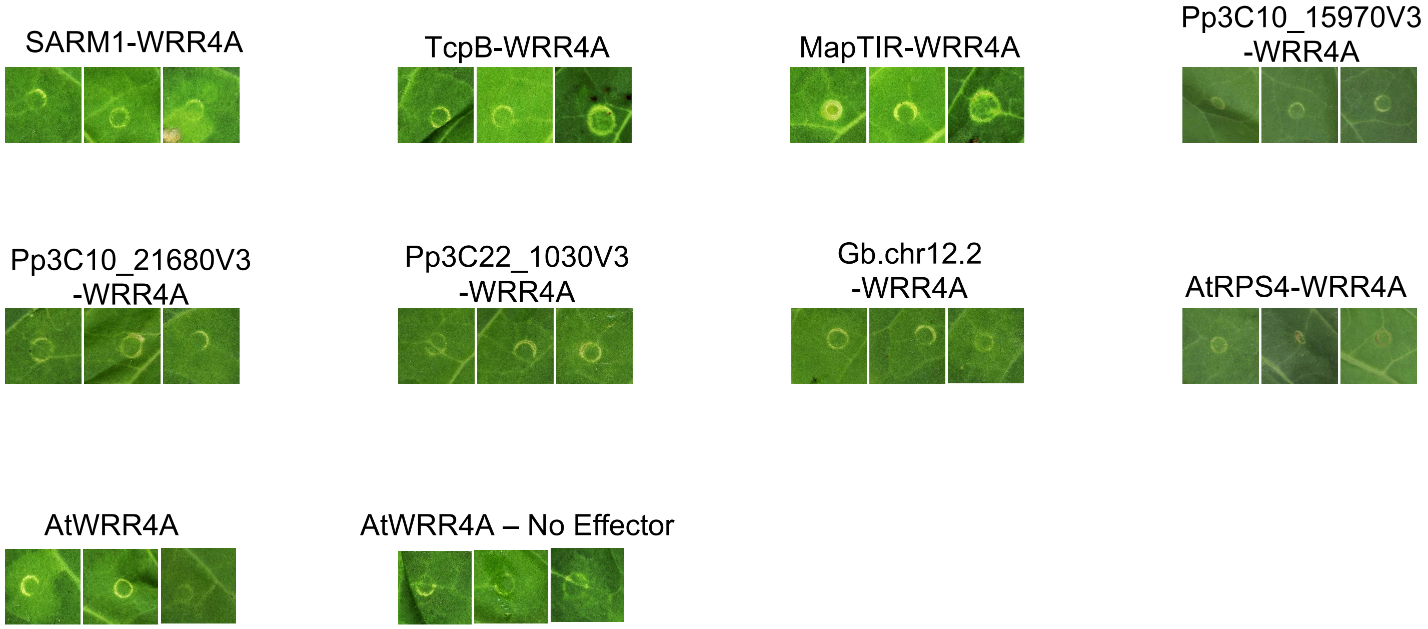


**Supplementary Dataset 2. Macroscopic Cell Death Phenotypes.**
**(J)** Representative cell death phenotypes for TIR-WRR4a fusions in *eds1 N. tabacum*, photographed 7 days post-infiltration, and cropped from independent leaves.
**(K)** Representative cell death phenotypes for TIR-WRR4a fusions in *RNAi::EDS1 N. tabacum*, photographed 7 days post-infiltration, and cropped from independent leaves.
