## Supplementary Data 2 for "Canonical NLR immune receptor architecture enforces EDS1-dependency onto divergent TIR domains"

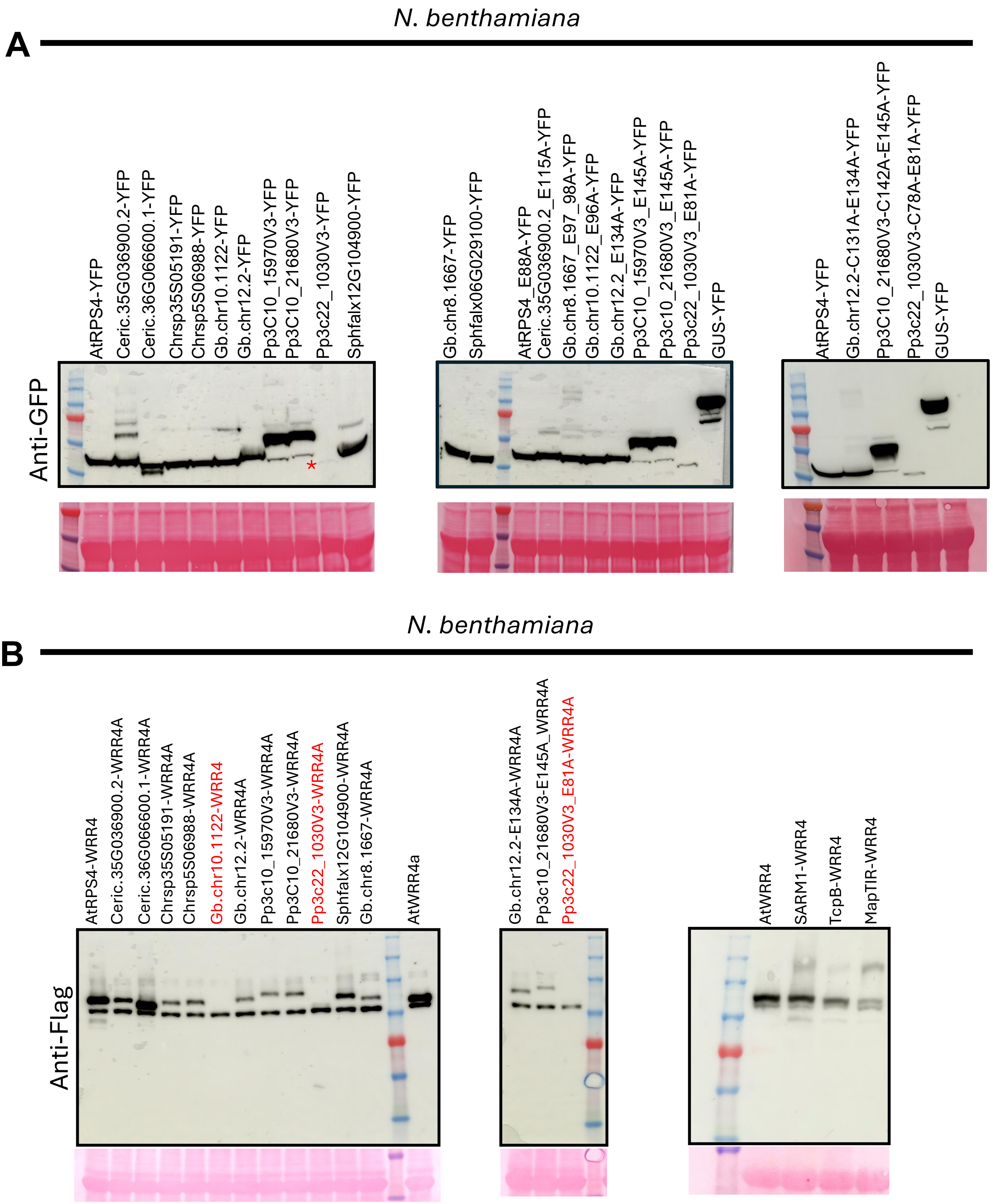


**Supplementary Dataset 1. Protein Immunoblotting**

**(A)** Anti-GFP immunoblots of TIR domains fused to YFP in *eds1 Nicotiana benthamiana*. Tissue was harvested 2 days post-infiltration for proteins inducing cell death and 3 days post-infiltration for proteins not inducing cell death. Weak bands are marked by a red asterisk.

**(B)** Anti-Flag immunoblots of TIR-WRR4A fusions in wild-type *N.benthamiana*. Tissue was collected 3 days post-infiltration. Red text indicates constructs for which no band was visible.

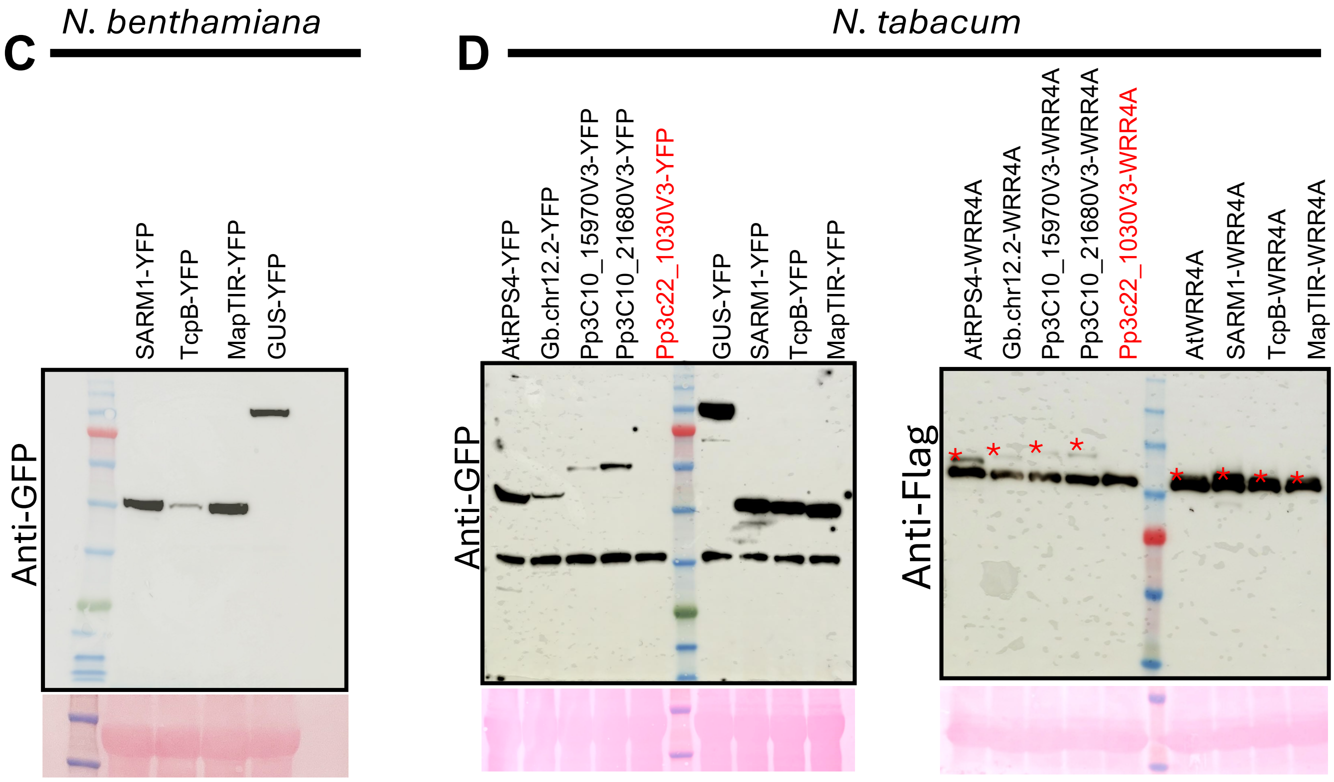


**Supplementary Dataset 1. Protein Immunoblotting**
**(C)** Anti-GFP immunoblots of animal and bacterial TIR-YFP fusions in *N.benthamiana*. Samples were collected 2 days post-infiltration.
**(D)** Anti-GFP and anti-Flag immunoblots for plant, animal and bacterial TIR-YFP and TIR-WRR4A fusions in *N.tabacum.* Samples were collected 2 days post-infiltration for proteins inducing cell death and 3 days post-infiltration for proteins not inducing cell death. Weak bands are marked with a red asterisk. Red text indicates constructs for which no band was visible.
